# The food–gut axis: the phyllosphere virome of fresh vegetables is diverse, abundant, and transmitted to the human gut virome

**DOI:** 10.1101/2024.09.09.612165

**Authors:** Ji-Woo Park, Yeo-Eun Yun, Jin Ah Cho, Su-In Yoon, Su-A In, Eun-Jin Park, Min-Soo Kim

## Abstract

Fresh vegetables harbor diverse microorganisms on leaf surfaces, yet their viral communities remain unexplored. We investigated the diversity and ecology of phyllosphere viromes of six leafy green vegetables using virus-like particle (VLP) enrichment and shotgun metagenome sequencing. On average, 9.2 × 10^7^ viruses were present per gram of leaf tissue. The majority (93.1 ± 6.2%) of these viruses were taxonomically unclassified. Virome compositions were distinct among vegetable types and exhibited temporal variations. Virulent phages with replication-enhancing auxiliary metabolic genes (AMGs) were more dominant than temperate phages with host fitness-benefiting AMGs. Analysis of 1,498 human fecal VLP metagenomes revealed that approximately 10% of vegetable viruses were present in the human gut virome, including viruses commonly observed in multiple studies. These gut-associated vegetable viruses were enriched with short-term vegetable intake, and depleted in individuals with metabolic and immunologic disorders. Overall, this study elucidates the ecological contribution of the fresh vegetable virome to human gut virome diversity.

## Introduction

The westernization of modern societies has brought about a decline in the biodiversity of the human gut microbiome^1,2^, in part because of reduced exposure to the natural environment^3,4^. Dietary plant fibers significantly influence the diversity and activity of the gut microbiome as these fibers serve as carbon and energy sources for fermentable bacteria^5^. Accordingly, frequent intake of plant-based foods helps to maintain the diversity of the gut microbiome in industrialized countries^6^. However, current research primarily emphasizes the impact of the dietary fiber content of these foods, overlooking the potential contribution of the microorganisms they carry.

Fresh vegetables harbor different types of microbial populations on their leaves^7^. The diversity of the vegetable phyllosphere microbiome is influenced by cultivation region^8,9^ and season^10^, agricultural practices^7^, and postharvest handling^11,12^. The phyllosphere microbiome improves crop quality during the preharvest stage^13^, while also influencing crop safety during the postharvest stage from production facilities to consumers^14^. Similar to microorganisms found in fermented foods^15,16^, those present on fresh vegetables can be transferred to the gut^17^ and may therefore influence human health. Although the importance of the vegetable microbiome is recognized, most research has focused on bacterial and fungal communities within vegetables.

Viruses, predominantly bacteriophages, are key members of microbial communities in natural ecosystems^18^. Viruses shape bacterial community composition through their lytic lifestyle^19^ and modulate the functional phenotype of bacterial hosts via auxiliary metabolic genes (AMGs)^20^. They also indirectly influence eukaryotic hosts through microbiome alterations^21,22^. Despite their ecological significance, characterizing viral communities (viromes) on fresh vegetables remains challenging, primarily because of difficulties in directly sequencing limited amounts of viral DNA^23^. Moreover, limitations in annotating viral sequences hinder understanding of the dynamics and functions of viromes, particularly in the human gut^24^. Therefore, investigating the fresh vegetable virome and its potential transmission to the gut is expected to elucidate the ecological origins of gut-resident viruses and their potential impact on health.

In this study, we investigated the phyllosphere virome of six leafy green vegetables (broccoli, spinach, perilla, red lettuce, green lettuce, and romaine lettuce) using a combination of virus-like particle (VLP) enrichment and shotgun metagenome deep-sequencing without prior random amplification. We characterized the total viral load, and the diversity, variability, and functional potential of the vegetable phyllosphere virome. Based on our *de novo* vegetable virome database, we explored the potential transfer of vegetable-associated viruses to the human gut by comparing their genomes with 1,498 fecal VLP metagenomes from 13 studies (huVLPs) and those in human gut virome databases (GVDs). The results of this study enhance our understanding of the viral ecology of fresh vegetables, and illuminate the role of the food-gut axis in shaping and maintaining the healthy human gut virome.

## Results

### Reconstruction of viral genomes from the VLP metagenomes of fresh vegetables

Leaf samples of six vegetables (n = 60) were collected from retail stores between 2019 and 2021 (Extended Data Fig. 1 and Supplementary Table 1). Leaf samples were homogenized to detach epiphytic and endophytic microorganisms including viruses, and then, VLPs were isolated. The metagenomic DNA of VLPs was deep-sequenced without random amplification, which avoids composition bias toward single-stranded circular viral genomes^23,25^. After removing plant-derived reads, an average (± s.d.) of 2,186,429 ± 3,724,274 high-quality paired-end reads per sample were obtained, accounting for 9.9 ± 15.0% of raw reads. Sixteen samples failed due to insufficient VLP-derived DNA for sequencing directly. The reads were assembled into 19,918 large contigs (≥2.5 kb). A combination of four *in silico* viral sequence identifications found 9,773 viral contigs, accounting for 33.8 ± 33.0% of the high-quality reads (Extended Data Fig. 2a). The completeness of viral contigs was estimated using CheckV^26^. One-fourth (23.8 ± 25.7%) of viral contigs had no amino acid similarities with previously known genomes. The remaining contigs were classified into low-quality (66.5 ± 23.0%), medium-quality (5.2 ± 6.5%), high-quality (2.9 ± 4.0%), and complete viral genomes (1.7 ± 2.5%). In total, the qualified contigs accounted for 78.3 ± 26.8% of the reads mapped to the viral contigs (Extended Data Fig. 2b), suggesting that a substantial number of vegetable viruses are unexplored. Finally, a virome dataset comprising 8,137 non-redundant viral genomes was established, which consisted of 198 circular (40,201 ± 28,855 bp) and 7,939 linear genomes (9,060 ± 13,088 bp). Using ViromeQC^27^, we quantified prokaryotic ribosomal RNA and single-copy marker genes in five datasets, each representing different stages of sequence processing: raw reads, preprocessed reads, high-quality reads, reads for large contigs, and reads for viral contigs. The viral enrichment score sharply increased from raw reads to reads for viral contigs (*P* < 0.0001) (Extended Data Fig. 2c), indicating that this study successfully obtained genomic sequences enriched in viral information from fresh vegetable samples.

### Fresh vegetables harbor an abundant and specific phyllosphere virome

We quantified the total viral abundance of vegetable viromes by adding *Escherichia coli* T4 phage to samples, the amount of which was pre-determined. Samples treated with T4 phage had an average of 2,587 ± 6,446 reads (0.2 ± 0.8%), whereas untreated samples had no T4 phage reads (*P* = 0.0003) (Extended Data Fig. 2d), validating the use of T4 phage as an internal standard for quantifying the total viral load of fresh vegetables. Based on the length-normalized read number of the T4 phage genome, an average of 9.8 × 10^7^ ± 2.2 × 10^8^ viral genomes were estimated per gram of leaf. The number of viral genomes was highest in spinach, lowest in perilla, and similar among the others (*P* < 0.05) (Fig. 1a). The number of viral genomes was significantly lower in pesticide-free samples than in conventional and organic samples (*P* < 0.05) (Fig. 1b). The number of viral genomes did not differ between sampling periods or distributors (Fig. 1c,d). These results suggest that numerous viruses are present on fresh vegetables, and that their abundance varies depending on vegetable type and farming method used. We also quantified total bacterial abundance and estimated an average of 3.1 × 10^7^ ± 1.3 × 10^8^ bacterial genomes per gram of leaf. Similar to viral genomes, the number of bacterial genomes was highest in spinach (Fig. 1e), and lowest in pesticide-free samples (*P* < 0.01) (Fig. 1f). The number of bacterial genomes did not differ between sampling periods or distributors (Fig. 1g,h). These similarities between the total viral and bacterial abundances suggest that the two are closely linked. Based on these observations, an average of 1.2 ± 12.7 virus-to-bacteria ratio (VBR) was determined across vegetables (Extended Data Fig. 2e). According to recent metagenome-based VBR estimations^28^, these values fall within the expected range for planktonic ecosystems, although the VBRs were variable. Considering that VBR roughly serves as an indicator of viral infection strategies in a given environment^29^, these results suggest that viral infection dynamics in the phyllosphere of fresh vegetables may be similar to those observed in relatively nutrient-poor environments^30^, wherein lytic dynamics dominate.

**Fig. 1.**
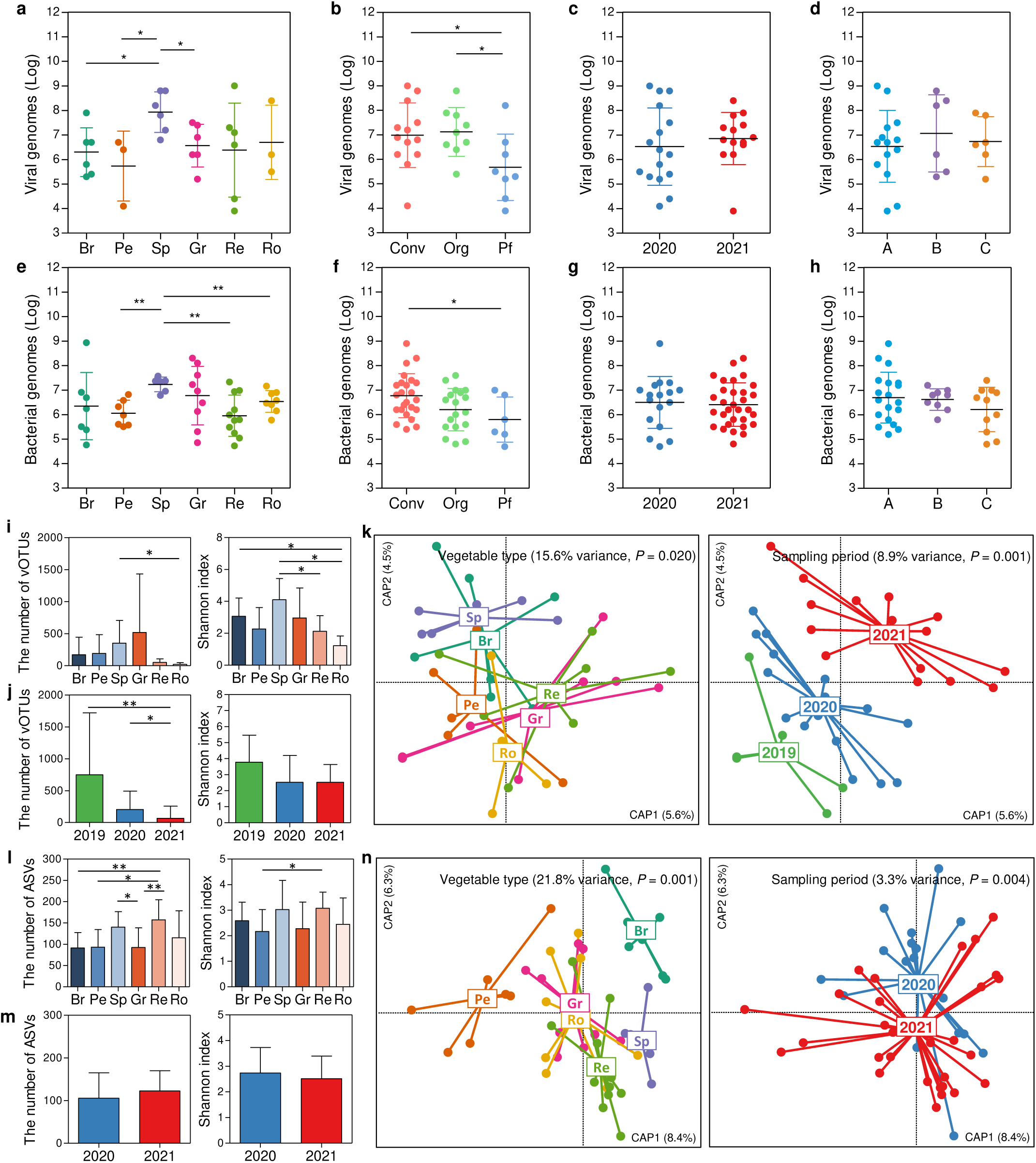
Variations in the total abundance, diversity, and composition of vegetable viromes and bacterial microbiomes by vegetable type. **a–d,** The number of viral genomes per gram of leaf among the vegetable types **(a)**, farming methods **(b)**, sampling periods **(c)**, and vendors **(d)**. **e–h,** The number of bacterial genomes per gram of leaf among the vegetable types **(e)**, farming methods **(f)**, sampling periods **(g)**, and vendors **(h)**. **i,j,** The number of observed vOTUs (left panel) and Shannon index (right panel) among the vegetable types **(i)** and sampling periods **(j)**. **k,** The best-fit model of viromes based on vegetable type (left panel) and sampling period (right panel) from distance-based redundancy analysis with Bray-Curtis dissimilarity. **l,m,** The number of observed ASVs (left panel) and Shannon index (right panel) among the vegetable types **(l)** and sampling periods **(m)**. **n,** The best-fit model of bacterial microbiomes based on vegetable type (left panel) and sampling period (right panel) from distance-based redundancy analysis with Bray-Curtis dissimilarity. Abbreviations: Br, broccoli; Pe, perilla; Sp, spinach; Gr, green lettuce; Re, red lettuce; Ro, romaine lettuce; Conv, conventional; Org, organic; and Pf, pesticide-free. All data are presented as mean ± S.D. Statistical significance was determined by two-tailed Mann-Whitney test. Symbol: *, *P*-value < 0.05; and **, *P*-value < 0.01.

The viral genomes were clustered into species-level taxonomic groups^31^, resulting in a total of 6,363 virus operational taxonomic units (vOTUs; 239 ± 485 vOTUs) (Supplementary Table 2). Significant differences were observed in the number of vOTUs and/or the Shannon index among the vegetable types and sampling periods (*P* < 0.05) (Fig. 1i,j), but not among the farming methods or distributors (Extended Data Fig. 3a,b). Virome composition was compared according to four host and environmental factors (vegetable type, sampling period, farming method, and distributor) using distance-based redundancy analysis (db-RDA). The best-fit model was constrained to vegetable type and sampling period, which explained 15.6% and 8.9% of total variance, respectively (*P* < 0.05) (Fig. 1k). Specifically, the viromes of the three lettuce types were separated from those of non-lettuces along the first axis of the db-RDA (*P* = 0.020), and the viromes of the three sampling periods were separated along the second axis (*P* = 0.001). These results suggest that the vegetable virome is host-specific, and changes temporally.

Similar patterns were observed in the bacterial microbiome. Based on 1,154 amplicon sequence variants (ASVs; 116 ± 52 ASVs), significant differences were observed in the number of ASVs and the Shannon index among vegetable types (*P* < 0.05), but not among sampling periods, farming methods, or distributors (Fig. 1l,m and Extended Data Fig. 3c,d). The best db-RDA model fitted by vegetable type and sampling period was selected, which explained 21.8% and 3.3% of the total variance, respectively (*P* < 0.05) (Fig. 1n). The bacterial microbiomes of the three lettuces were distinct from those of non-lettuce vegetables along the first axis (*P* = 0.001), and the bacterial microbiomes of the three sampling periods were separated along the second axis (*P* = 0.004). Therefore, strong co-variation between viral and bacterial microbiomes suggests that the ecology of the virome is closely associated with the bacterial microbiome in the vegetable phyllosphere.

### Fresh vegetable viruses are largely uncharacterized

We next determined the taxonomic diversity of the vegetable virome according to the International Committee on Taxonomy of Viruses taxonomy^32^. A gene-sharing network was constructed with the viral genomes of the 2,003 vOTUs (31.5%), the NCBI RefSeq Viral Genome, and IMG/VR using vConTACT2^33^, resulting in a total of 1,062 viral clusters (VCs; 65 ± 108 VCs) representing genus-level taxonomic groups (Fig. 2a). Of those, 107 vOTUs (5.3%) in 64 VCs were classified into 49 viral genera of nine families, and 433 vOTUs in 147 VCs were assigned to 10 families. As a result, 5,823 vOTUs (93.1 ± 6.3%) could not be classified into any genus or family (Fig. 2b), underscoring that viruses present on fresh vegetables are largely uncharacterized.

**Fig. 2.**
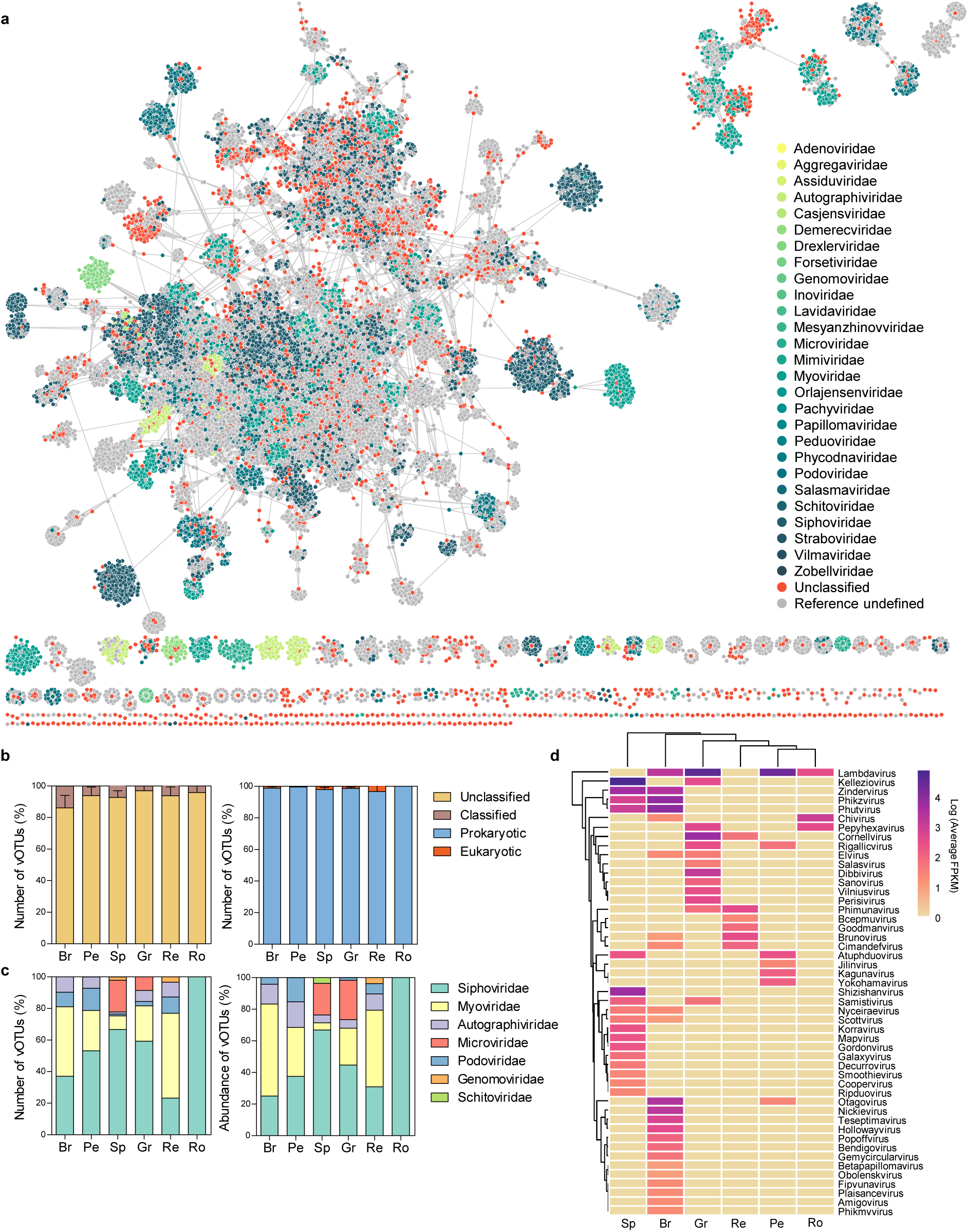
Taxonomic diversity of fresh vegetable viromes. **a,** A gene-sharing network was constructed with the genomes of vOTUs, IMG/VR and NCBI Viral RefSeq. The taxonomy of viral clusters was colored according to their viral families. **b,** The percentage of taxonomically classified vOTUs (left panel) and the percentage of prokaryotic vOTUs among the classified vOTUs (right panel). **c,** the percentage of viral families (left panel) and their abundance (right panel) in six vegetables. Only viral families with an average percentage ≥ 1% are shown. **d,** The composition of viral genera in six vegetables. Hierarchical clustering was performed using the average method.

Taxonomically identified vOTUs were mostly classified as prokaryotic viruses (98.4 ± 3.6%) (Fig. 2b), with most assigned to four families within the *Caudovirales* order: *Siphoviridae* (50.4 ± 38.1%), *Myoviridae* (29.2 ± 30.1%), *Podoviridae* (6.9 ± 9.5%), and *Autographiviridae* (6.5 ± 9.7%) (Fig. 2c). Viral abundance was largest in the *Siphoviridae* family (44.8 ± 42.3%), followed by the *Myoviridae* (31.9 ± 38.0%) and *Autographiviridae* families (9.1 ± 15.6%) (Fig. 2c). Single-stranded *Microviridae* vOTUs were observed only in spinach (20.0 ± 44.7%) and green lettuce (24.6 ± 28.7%). At the genus-level, 31 genera (63.3%) were observed in only a single vegetable type, while *Lambdavirus* was observed across four vegetables (Fig. 2d). These results suggest that host-specific bacteriophages dominate the phyllosphere virome of fresh vegetables.

### Virulent and temperate phages differently interact with their bacterial hosts

To understand how vegetable viruses interact with their hosts, we focused on viral lifestyles by searching for lysogeny-associated genes and host regions flanking viral genomes. A total of 484 vOTUs were identified as temperate phages (8.5 ± 7.4% of the vOTUs and 17.5 ± 23.7% of the total abundance) (Fig. 3a), indicating that virulent phages predominate the vegetable virome, consistent with the VBR prediction in Extended Data Fig. 2e. This fraction remained relatively consistent regardless of genome completeness and sampling periods (Extended Data Fig. 4a,b). Next, we predicted prokaryotic hosts of virulent and temperate vOTUs using iPHoP^34^, combined with sequence matches of CRISPR spacers and tRNAs. A total 1,059 vOTUs (16.6%) were linked to a single bacterial host. For both phage types, hosts most frequently belonged to *Gammaproteobacteria* (59.1 ± 32.5% of virulent vOTUs; 40.3 ± 34.0% of temperate vOTUs), followed by *Actinomycetes* (13.9 ± 22.8%; 20.0 ± 28.2%) and *Alphaproteobacteria* (8.4 ± 13.9%; 36.2 ± 37.8%) (Extended Data Fig. 4c,d). However, the bacterial phyla, families, and genera infected with virulent vOTUs were significantly more diverse than those infected by temperate vOTUs (*P* < 0.05) (Fig. 3b), although this pattern varied depending on bacterial taxa (Fig. 3c). These results suggest that on vegetables, virulent phages are more common and interact more broadly with bacterial hosts than temperate phages.

**Fig. 3.**
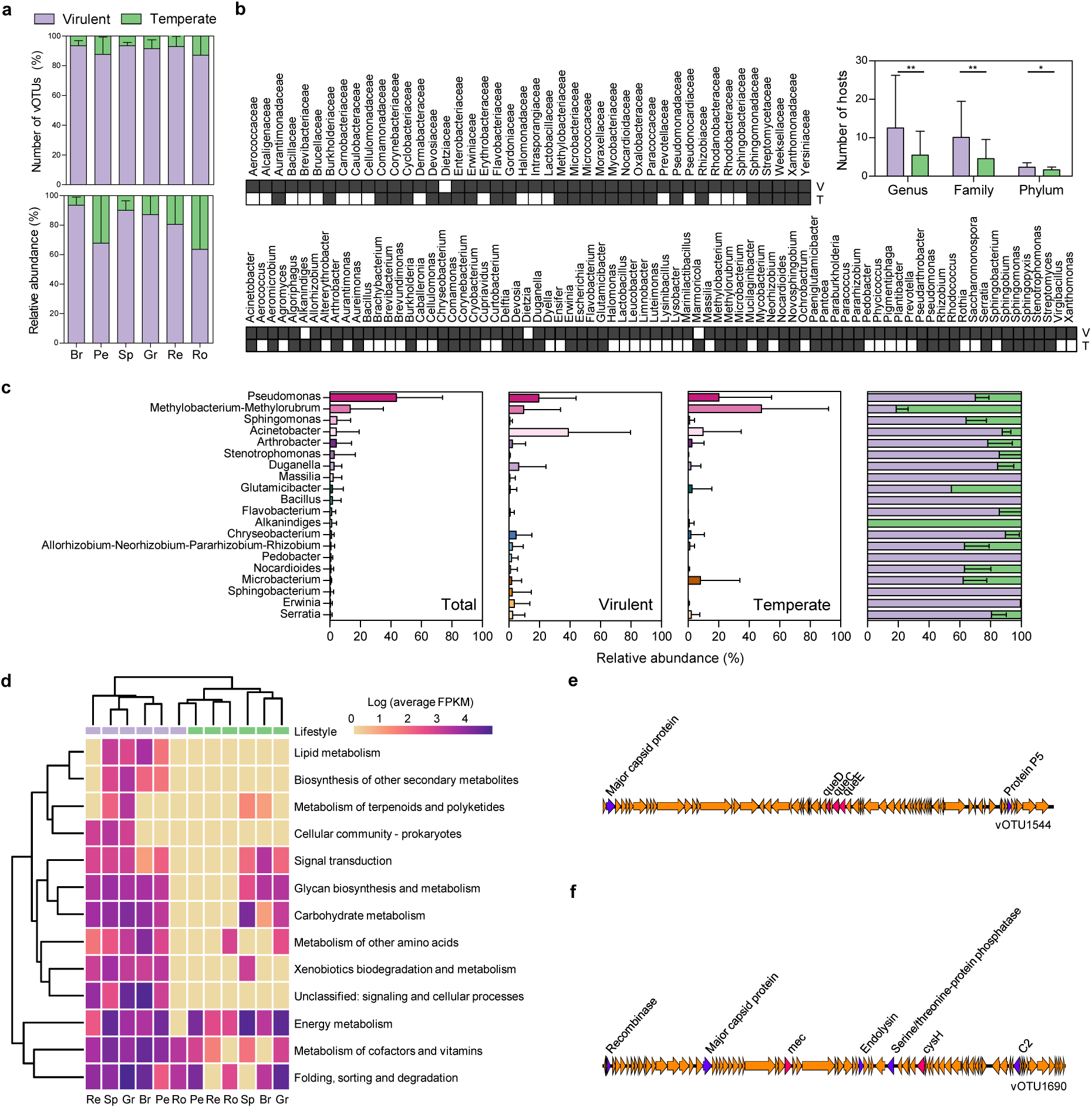
Viral lifestyle and AMG composition in the virome of fresh vegetables. **a,** The percentage of virulent and temperate vOTUs (upper panel), and their abundance (lower panel) among the vegetable types. **b,** Heatmap shows the bacterial hosts of virulent (V) and temperate (T) vOTUs at the family- (upper panel) and genus-level (lower panel). Bar chart shows comparisons between the number of bacterial hosts for virulent and temperate vOTUs at the phylum-, family-, and genus-level. **c,** The abundance of top 20 bacterial genera in bacterial communities (Total), virulent vOTUs (Virulent), and temperate vOTUs (Temperate) was shown. **d,** The composition of AMGs functions encoded in virulent and temperate vOTUs is shown among the vegetable types. Hierarchical clustering was performed using the average method. **e,f,** The genome map of virulent vOTU1544 **(e)** and temperate vOTU1690 **(f)** with their AMGs.

Next, we identified AMGs encoded in the genomes of virulent and temperate vOTUs using DRAM-v^35^ and VIBRANT^36^. A total of 458 AMGs were found in 345 virulent vOTUs, and 66 in 56 temperate vOTUs (Supplementary Table 3). DRAM-v identified the *phoH* (phosphate starvation-inducible) gene as the most common (36 AMGs), followed by AMGs related to xenobiotics biodegradation and metabolism (17) and glycan metabolism (9). VIBRANT identified AMGs involved in the metabolism of cofactors and vitamins (190), carbohydrate metabolism (190), and glycan metabolism (54) as the most common. Notably, the overall AMG composition differed significantly between virulent and temperate vOTUs (*P* = 0.012) (Fig. 3d). Specifically, AMGs involved in lipid metabolism, biosynthesis of other secondary metabolites, cellular community, and unclassified signaling and cellular processes were exclusive to virulent vOTUs (Fig. 3d). In addition, AMGs related to the metabolism of cofactors and vitamins, xenobiotics biodegradation and metabolism, and folding, sorting and degradation were enriched in virulent vOTUs, while energy metabolism AMGs were enriched in temperate vOTUs (*P* < 0.05) (Extended Data Fig. 4e). These findings align with the observations of Luo *et al*.^20^ describing that virulent viruses modulate translation efficiency for rapid replication, while temperate viruses adjust host metabolism to provide competitive advantages to their hosts. For example, virulent vOTU1544 encodes *queC*, *queD*, and *queE* genes for the synthesis of 7-cyano-7-deazaguanine, potentially influencing translation efficiency^37^ (Fig. 3e). By contrast, temperate vOTU1406 encodes *cysH* and *mec* genes, involved in assmilatory sulfate reduction^38^ (Fig. 3f). Collectively, these results indicate that virulent and temperate viruses interact differently with their hosts in the microbiome of fresh vegetables.

### Fresh vegetable viruses are transferred to the human gut

Based on environmental source information provided by IMG/VR, we found that the vegetable virome (927 vOTUs in 552 VCs) was associated with various ecosystems, including plant- (25.0 ± 13.7%) and soil-associated (22.7 ± 18.0%) ecosystems (Extended Data Fig. 5). Of these, we focused on 108 vOTUs (8.7 ± 19.0%) that were associated with viral genomes derived from human-associated ecosystem. Zhang *et al*.^39^ demonstrated that the intestinal RNA virome largely consists of plant-infecting viruses derived from dietary plants, a finding corroborated by later studies^40,41^. We hypothesized that similar observations could be made for the DNA gut virome. We compared vOTU genomes with those in IMG/VR and three GVDs (Gut Phage Database^42^, human Gut Virome Database^43^, and Metagenomic Gut Virus^44^), and identified 61 vOTUs that were associated with 92 GVD genomes at the genus or species level (Extended Data Fig. 6a). Particularly, five vOTUs showed species-level genomic similarities with 24 GVD genomes (Supplementary Table 4), and vOTU1690 had an identical genome to gvd30371 (Extended Data Fig. 6b).

Typically, only a small fraction of foodborne microorganisms is found in the human gut microbiome^17,40^. GVDs are predominantly constructed from bulk metagenomes^45^, which restricts in-depth viral detection, compared with VLP metagenomes^46,47^. Based on these considerations, we tested our hypothesis using virome datasets generated through VLP enrichment, which was expected to allow more sensitive detection of vegetable-derived viruses. We collected 1,498 VLP metagenomes from 1,306 individuals across 13 studies, representing diverse demographics and health conditions (Fig. 4a and Supplementary Table 5). High-quality reads were compared with vOTU genomes. crAssphage genome was used as a control because of its high abundance and prevalence in the human gut virome^48^. We identified 627 vOTUs (25 ± 40 vOTUs) as gut-associated (Fig. 4b and Supplementary Table 6). An average of 130 ± 97 vOTUs were detected per study, with low abundance (0.6 ± 1.6 fragments per kilobase million, FPKM, per vOTU and 41.4 ± 89.8 FPKM for total vOTUs) but high prevalence (97.8%), compared with crAssphage (109.6 ± 382.9 FPKM and 77.2%, respectively). While this number appears to reach a plateau within huVLPs (Fig. 4c), it is expected to increase as additional studies are conducted (Fig. 4d). Of those, 101 vOTUs were observed in more than three studies, and 46 vOTUs were observed in more than four studies (Fig. 4e). These global vOTUs were derived from all vegetables, and were repeatedly observed in one-year longitudinal samples^49^ (Extended Data Fig. 7). For bacterial hosts, six global vOTUs were identified as *Pseudomonas*, four as *Escherichia*, three as *Pedobacter*, two as *Chryseobacterium*, and one each as *Pantoea*, *Duganella*, *Burkholderia*, *Acinetobacter*, and *Erwinia*. These genera have been reported in human gut metagenome-assembled genome datasets^50,51^, and were recently identified as vegetable-derived intestinal bacteria^17^. However, these results were obtained from eight huVLP datasets, with only a few vOTUs (12 ± 7 vOTUs per study) detected in five of the datasets. To further investigate this, we classified huVLPs into three categories based on sequencing depth: high (≥3 million reads), medium (1–3 million reads), and low (<1 million reads). The five datasets with a significantly lower number of vOTUs belonged to the low- or medium-depth group (*P* < 0.05) (Fig. 4f), indicating the need for deep-sequencing to effectively detect vegetable viruses in VLP metagenomes. Collectively, these findings suggest that viruses from fresh vegetables are transferred to the human gut, albeit as a relatively small proportion of the gut virome.

**Fig. 4.**
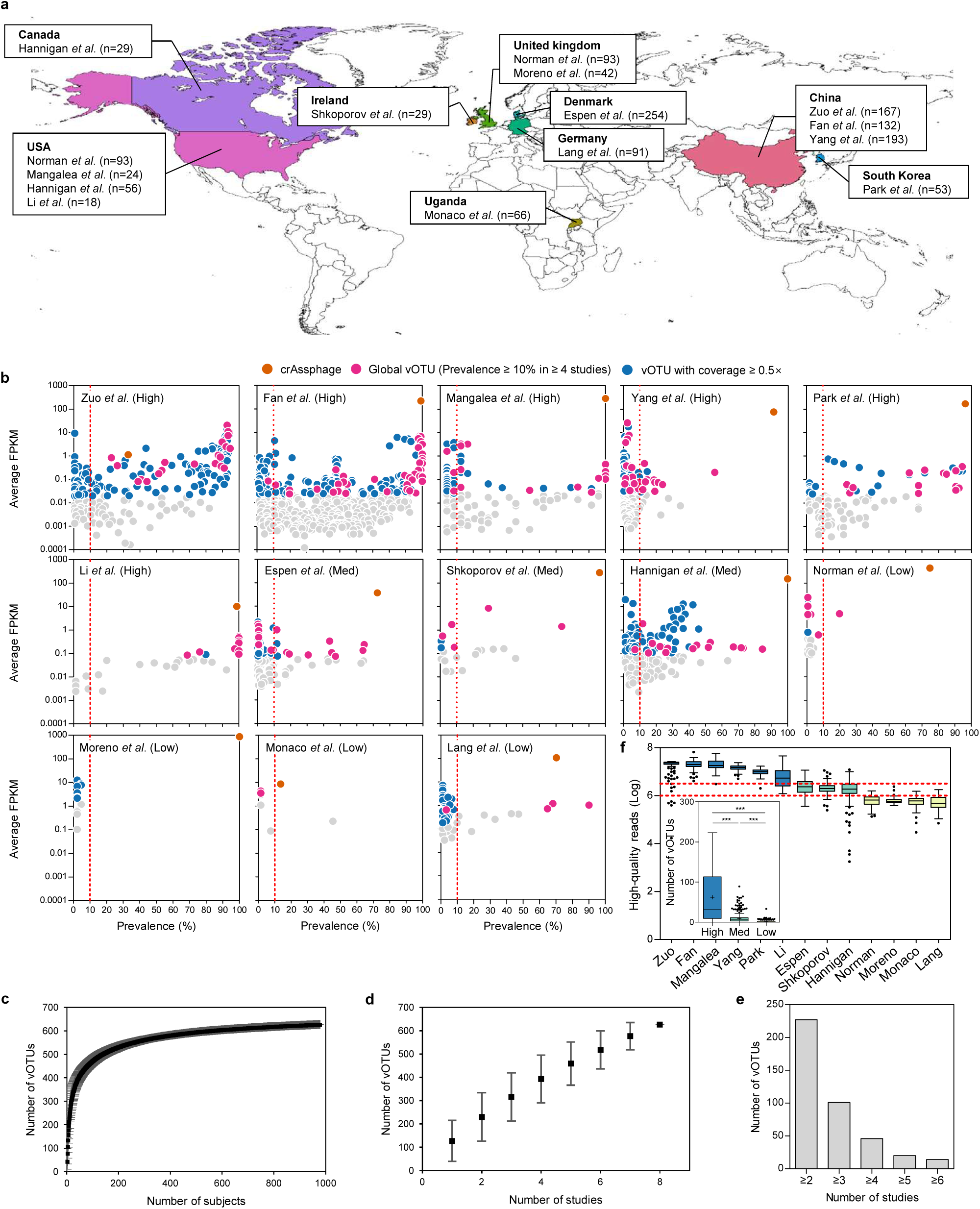
Abundance and prevalence of gut-associated vegetable viruses in the human gut virome. **a,** The geography of human gut VLP metagenomes collected for this study. **b,** The abundance and prevalence of gut-associated vOTUs in the human gut VLP metagenomes. Orange dots represent crAssphage, pink dots represent global vOTUs, and blue dots represent vOTUs with ≥0.5× coverage. **c,d,** Accumulation curves for vOTUs based on subject **(c)** and study **(d)**. **e,** The number of global vOTUs shared across different numbers of studies. **f,** The number of high-quality reads across the human gut VLP metagenomes. Box plots are displayed using Tukey’s method. Statistical significance was determined by two-tailed Mann-Whitney test. Symbol: ***, *P*-value < 0.001.

### Short-term vegetable intake induces viral transmission from fresh vegetables

We examined whether viral transmission from fresh vegetables occurs through the consumption of these vegetables. We recruited 53 healthy individuals (Supplementary Table 7) and analyzed their fecal virome based on short-term and long-term dietary patterns, characterized by intake amounts of 16 and 20 food groups, respectively. The composition of 66 gut-associated vOTUs varied significantly with the short-term intake of two food groups: vegetables and fruits (*P* < 0.05) (Fig. 5a). This was confirmed by redundancy analysis, which revealed that variations in the vOTUs were explained the most by four food groups in the following order: vegetables (*r*^2^ = 0.0357), potatoes (*r*^2^ = 0.0351), legumes (*r*^2^ = 0.0245), and fruits (*r*^2^ = 0.0230) (*P* < 0.05) (Fig. 5b). Specifically, 24 vOTUs showed significant enrichment or depletion with short-term intake of the four groups, with the largest positive associations observed for vegetable (*P* < 0.05) (Fig. 5c). Subjects with a high vegetable intake had significantly more vOTUs than those with low or medium consumption (*P* < 0.05) (Fig. 5b). These results provide evidence that frequent, short-term consumption of dietary vegetables may facilitate the transmission of viruses from fresh vegetables. These results were not replicated in long-term dietary patterns. This may also explain why the majority of vOTUs (49.2 ± 25.2%) had a prevalence of less than 10% among subjects in each study. However, the abundance of vOTUs in huVLPs did not correlate positively with their abundance and prevalence in the original samples (Extended Data Fig. 8a,b), suggesting that high vegetable consumption does not fully facilitate the equal transmission of all viruses to the gut. This is further reflected in the composition comparisons between bacterial genera in the vegetable virome and bacterial hosts in huVLPs. Indeed, only a subset of abundant vOTUs appears to be frequently transferred to the human gut (Fig. 5d). This cannot be explained by viral lifestyle as well (Extended Data Fig. 8c). These findings allow speculation that viral transmission from fresh vegetables may be selective process.

**Fig. 5.**
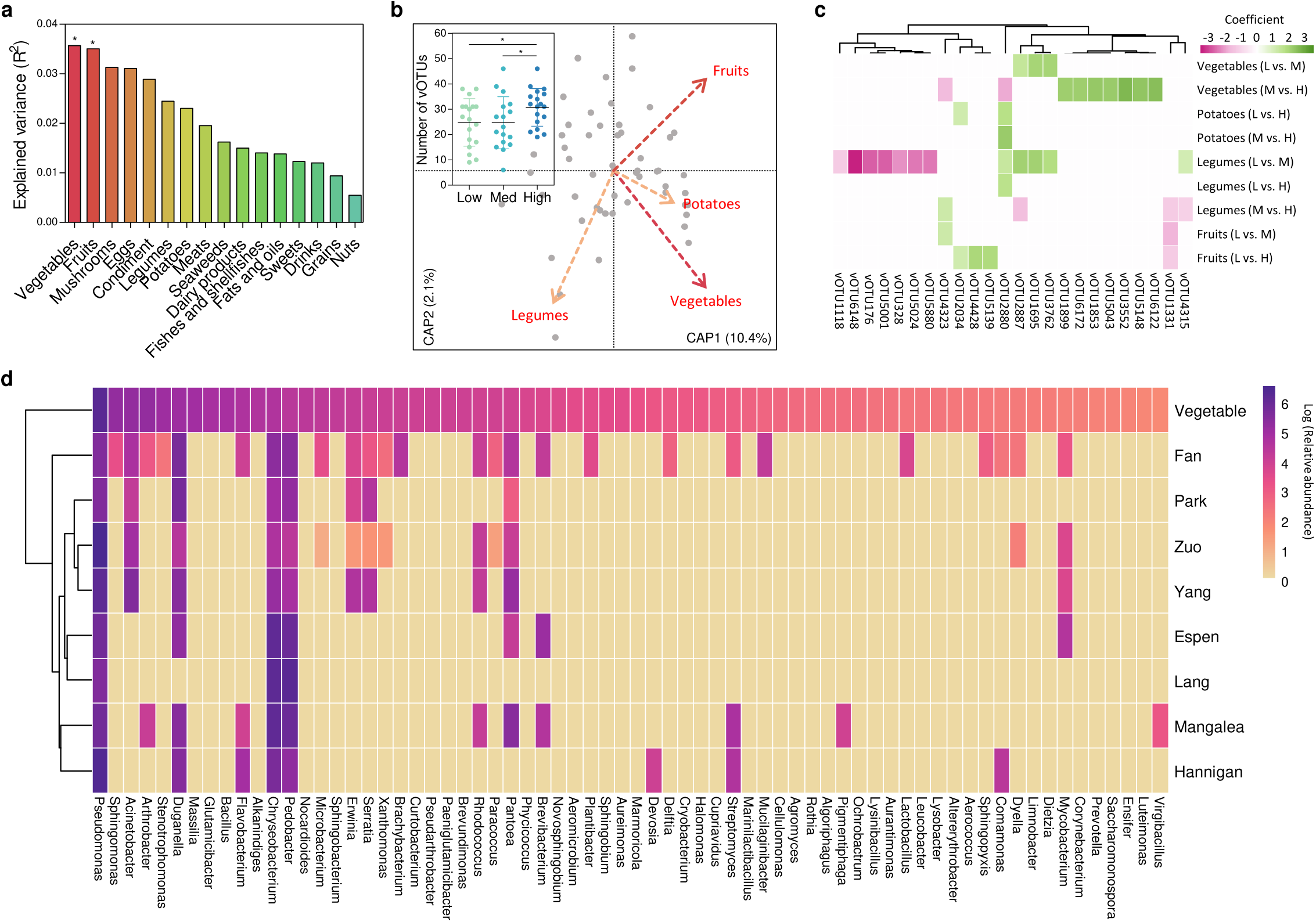
Correlations between the composition of gut-associated vOTUs and short-term dietary patterns. **a,** The variances of gut-associated vOTU composition explained by the 16 food group of short-term dietary patterns. **b,** The best-fit model of gut-associated vOTUs based on four food groups (Vegetables, Fruits, Potatoes, and Legumes) from distance-based redundancy analysis with Jaccard distance. Box plot shows the number of gut-associated vOTUs among the high-, medium-, and low-amount consumption of vegetable. **c,** Gut-associated vOTUs significantly enriched or depleted in groups with high-, medium-, and low-amount vegetable consumption. **d,** The composition comparisons between bacterial genera in the vegetable virome and bacterial hosts in VLP metagenomes. Hierarchical clustering was performed using the average method. Abbreviations: H, high-; M, medium-; and L, low-amount vegetable consumption.

### Vegetable viruses are reduced in individuals with metabolic and inflammatory disorders

Disease-specific alterations in the intestinal virome (referred to as “virome dysbiosis”) have been observed in individuals with metabolic and inflammatory disorders^24,52–56^. In this context, we characterized the dynamics of gut-associated vegetable viruses in virome dysbiosis associated with obesity^55^, ulcerative colitis^53^, colorectal cancer/adenoma^52^, alcoholic liver disease (ALD)^54^, and type 2 diabetes (T2D)^56^. The composition of gut-associated vOTUs in the disease groups differed significantly from that in healthy controls across all conditions (*P* < 0.01) (Fig. 6a). Specifically, viral diversity (Shannon index) and/or relative abundance (FPKM) of the vOTUs was significantly lower in the disease groups than in healthy controls (*P* < 0.05) (Fig. 6a). Also, the abundance of most vOTUs was significantly lower in the disease groups than in healthy controls (*P* < 0.05) (Fig. 6b). These findings suggest that a reduction in vegetable-derived viruses is a common feature of the virome dysbiosis observed in metabolic and inflammatory disorders.

**Fig. 6.**
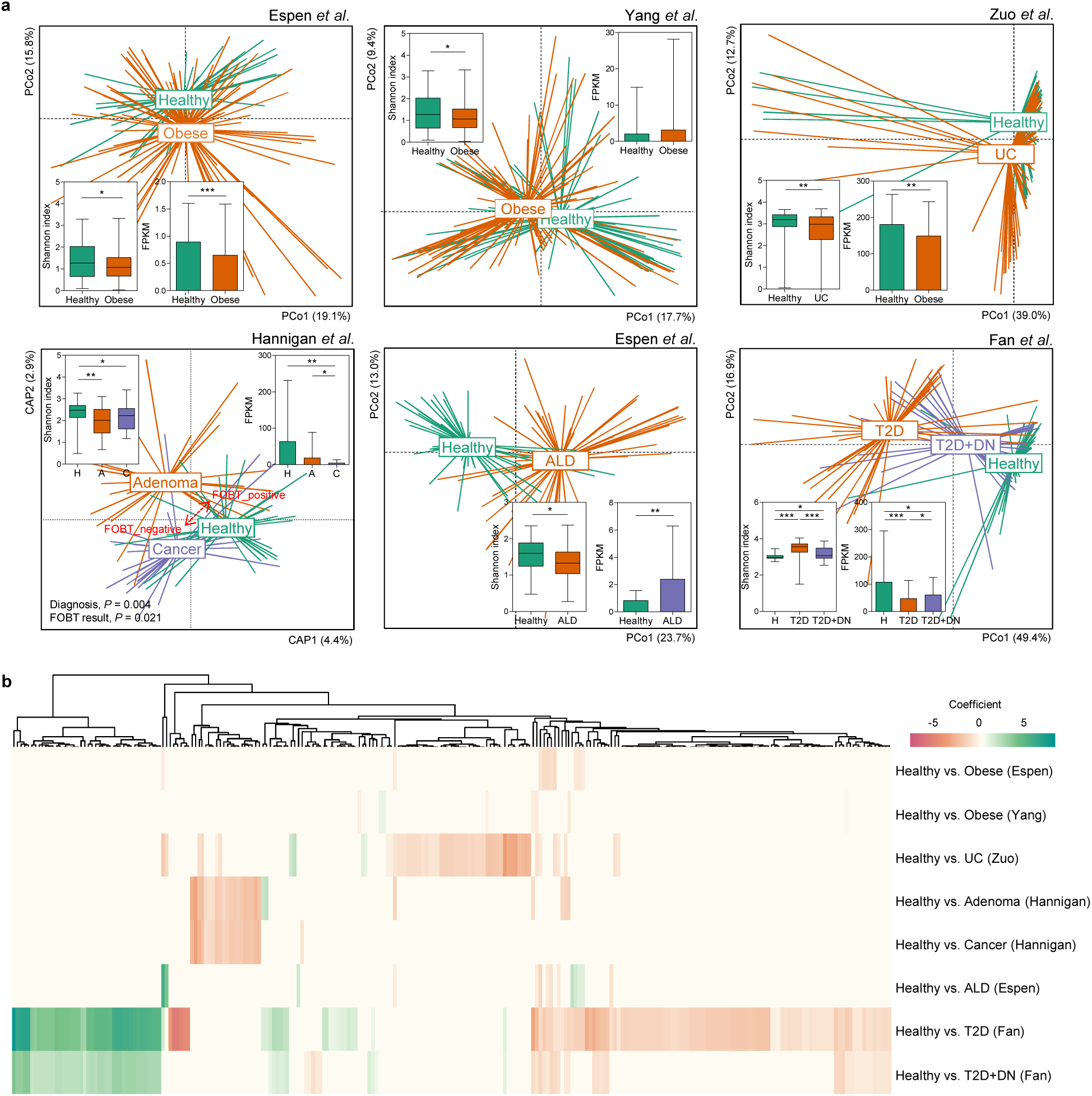
Alterations in the diversity and composition of gut-associated vegetable viruses in with metabolic and inflammatory disorders. **a,** The comparisons of gut-associated vOTU composition, Shannon index, and total FPKM between healthy and disease groups are shown for obesity, ulcerative colitis (UC), colorectal cancer, alcoholic liver disease (ALD), and type 2 diabetes (T2D) with/without diabetic nephropathy (DN). The analyses were performed using principal coordinate analysis or distance-based redundancy analysis with Bray-Curtis dissimilarity. **b,** vOTUs significantly enriched or depleted in either healthy or disease group are shown for six datasets.

## Discussion

The consumption of fresh vegetables benefits human health through multiple mechanisms, including the provision of bioactive compounds^57^ and dietary fiber content, as well as the resulting microbial fermentation^58^. While the potential influence of microorganisms from fresh vegetables would also be expected, evidence for the microbiome-food-gut axis has been largely restricted to lactic acid-producing bacteria in fermented foods^15,16^. To explore this, the transmission of microorganisms from fresh vegetables to the host, and the interaction of microorganisms with the host’s gut microbiome requires elucidation. This study identified a high diversity, specificity, and variability of the phyllosphere viromes of fresh vegetables, as well as a virome-food-gut axis, revealing that fresh vegetables contribute significantly to the diversity and dynamics of the human gut virome.

The human gut virome is predominantly composed of DNA bacteriophages, with RNA viruses comprising a smaller proportion^41,49,59^. While RNA gut viruses are known to be primarily plant pathogenic viruses introduced through the consumption of plant foods including vegetables^39^, DNA gut viruses remain largely “viral dark matter” because of limited viral sequence information representing diverse environments, which hinders the identification of a healthy gut virome^60^. Our previous study suggested that temperate phages induced from lysogenic bacteria contribute to the composition of the DNA gut virome^61^, a finding confirmed by research on the infant gut virome^62^ and gut-derived bacterial prophages^63^. In this context, this study advances our understanding of human gut virome composition by demonstrating the horizontal virome transfer from fresh vegetables to the human gut. Specifically, on average, 10.0% of the vegetable virome was detected in global huVLP datasets, albeit at low abundance. This proportion is expected to increase as more viral genomes from fresh vegetables and the human gut virome are analyzed. Dozens of gut-associated vOTUs were shared across multiple, independent huVLP studies, each representing different regions, countries, and ethnicities. Although their abundance varied, their presence remained consistent within the same individuals over extended periods. Additionally, bacterial hosts associated with gut-associated vOTUs support the hypothesis of viral transmission from fresh vegetables, as these bacterial genera have been previously identified in human gut microbiomes^50,51^. All these findings provide evidence that viruses from fresh vegetables reach the distal colon, which aligns well with the recent study of Wicaksono *et al*.^17^ demonstrating the presence of fruit and vegetable-derived bacteria in the human gut. Collectively, DNA viruses from fresh vegetables may be an integral component of the human gut virome, and fresh vegetables are an environmental reservoir for human gut DNA viruses, similar to the RNA gut virus^39^.

Positive associations between gut-associated vOTUs and vegetable intake, both in terms of viral richness and composition, further support the hypothesis that viral transmission may occur through the consumption of fresh vegetables. These associations were observed only with short-term dietary patterns, rather than long-term patterns, suggesting that vegetable-associated viruses observed in the gut virome are likely acquired exogenously and transiently, rather than being part of a stable gut virome. This hypothesis is supported by high individuality in the abundance and prevalence of gut-associated vOTUs, which exhibited log-normal-like distributions in the huVLP datasets. Such distributions are similar to those observed for exogenously acquired microorganisms^64^, providing additional evidence for their transient nature in the gut ecosystem. The findings of Carmody *et al*.^65^ that raw plant foods disrupt gut microbiome physiology support this concept to be more feasible. Considering that most viral genera were specific to a particular vegetable type, consuming a variety of vegetables may contribute to a diverse individual gut virome^66^. However, viral transmission from fresh vegetables may not operate equally for all viruses, which is supported by the lack of associations between the prevalence and abundance of vOTUs on vegetables and those in the human gut. For example, *Pseudomonas*, *Escherichia*, *Chryseobacterium*, and *Pedobacter* were identified as hosts for global vOTUs, and are all known to possess mechanisms that resist colonization resistance by forming biofilms^67–69^ or interacting with the mucus layer^70^, allowing them to persist in the human gut. Moreover, the AMGs of gut-associated vOTUs, such as those encoding polygalacturonase for dietary pectin degradation, likely expand the metabolic niches of their hosts by enhancing nutrient utilization^71^. These characteristics suggest a model of viral transmission, wherein favorable viruses from fresh vegetables are selectively transmitted to the gut. Thus, transfer of viruses along with their bacterial hosts may be linked to the transmission and persistence of vegetable viruses in the gut.

Interestingly, a community of gut-associated vOTUs differed between individuals with metabolic and inflammatory disorders and healthy controls. These differences were characterized by a reduction in viral diversity and/or their relative abundance. Virome dysbiosis typically involves a reduction in overall viral diversity^52,53,55,56,72^, driven by the induction of temperate *Caudovirales* phages^73^, suggesting that the expansion of dysbiosis-associated phages may reduce the fraction of vegetable-derived viruses, potentially making them less detectable in sequencing data. Accordingly, our findings propose that gut-associated vegetable viruses could constitute a significant component of a healthy gut virome^60^ and may therefore be beneficial, rather than deleterious. While this behavior differs from the patterns observed in ectopic colonization by translocated oral bacteria in the intestine^74^, the distinct behavior of vegetable-derived viruses in the gut virome could serve as a biomarker for the dysbiotic state of the gut virome. Intriguingly, not all datasets such as the ALD^41^ or T2D^56^ studies exhibited reductions in viral diversity and abundance. Therefore, further validation of the “marker” hypothesis for gut-associated vegetable viruses is required through well-controlled experiments.

Based on our findings, consuming a typical serving of 85 grams of salad daily^75^ results in the ingestion of approximately 7.8 billion viruses, a number that can be increase by prophage induction caused by sauces or sweeteners^76,77^. Prolonged exposure to these viruses could influence the composition and function of the gut microbiome. For example, Hsu *et al*.^19^ demonstrated that even a single foreign phage can disrupt the composition of the gut microbiome and its associated metabolites in a mouse model. While vegetable-derived viruses were present in low amounts and were short-lived in the gut, their ecological behavior appears similar to that of foodborne probiotics^78^, suggesting that they might have comparable effects on the gut ecosystem, regardless of their abundance^79^. Given that vegetable-derived viruses are integral to a healthy gut virome, they may play a role in modulating the microbiome and subsequent phenotypic changes in both human subjects with metabolic syndrome^80^ and chemical-induced colitis mice^81^, following fecal virome transplantation from healthy donors. These observations indicate that frequent exposure to vegetable-derived viruses could have transient, individualized impacts on the assembly and dynamics of the gut microbiome and, consequently, on human health.

Our findings highlight the phyllosphere of vegetables as an overlooked part of the environmental virome and suggest that dietary vegetables act as vectors for the transmission of plant microbiome-associated bacteriophages to the human gut. Our *de novo* vegetable virome data represents only a fraction of the actual virome present on fresh vegetables. Expanding the collection of viral genomes from a wider variety of fresh vegetables, capturing actively infecting or integrated viruses from bulk metagenomes, addressing RNA viruses, and including preharvest samples could enhance the detection of vegetable-originated viruses in the gut virome. Further experimental investigations, combined with strain-level analysis of gut viromes and paired vegetable viromes, are necessary to corroborate our findings. Overall, this study significantly advances our understanding of the viral ecology of fresh vegetables and their interactions with the human gut, and the findings may also serve as a basis for future studies to investigate the impact of food-derived microorganisms on human health through the microbiome-food-gut axis.

## Methods

### Sample collection

Samples of broccoli (*Brassica oleracea var. italic*, *n* = 10), perilla (*Perilla frutescens var. crispa*, *n* = 9), spinach (*Spinacia oleracea*, *n* = 6), green leaf lettuce (*Lactuca sativa*, *n* = 11), red leaf lettuce (*Lactuca sativa*, *n* = 13), and romaine lettuce (*Lactuca sativa*, *n* = 11) were purchased from six retail stores located in Daejeon city and Pyeongtaek, Republic of Korea, during August 2019 to July 2021 (Supplementary Table 1). Samples labeled as conventional, pesticide-free, and organic were collected for different types of vegetables, according to their packaging labels. Samples were delivered and stored at 4 °C for no more than three days before further processing. Approximately 50-100 g of broccoli floret, 50-100 g of upper part of lettuce and perilla, and 10-30 g of a whole leaf of spinach were separated from the stem area using a sterile knife, and then cut into uniform pieces with a maximum width of 20 mm.

### Preparation of Escherichia coli T4 phage

*Escherichia coli* T4 phage DSM 4505 was prepared, according to Bonilla *et al*.^82^. Briefly, T4 phage was propagated by infecting *E. coli* DSM 613. Phage lysate was centrifuged at 4,000 ×g for 20 min at 4 °C, and the supernatant was filtered through 0.20-µm pore cellulose acetate syringe filters (Sartorius, Germany). The filtrate was mixed with 0.1 volume of chloroform (Sigma-Aldrich, USA) and incubated for 10 min at room temperature. The mixture was centrifuged at 4,000 ×g for 5 min at room temperature, and the supernatant was washed with 0.02-μm pre-filtered, sterilized sodium-magnesium (SM) buffer using Amicon^Ⓡ^ Ultra-15 Centrifugal Filters (100 kDa MWCO, Sigma-Aldrich, USA). The phage concentrate was serially diluted, filtered through 0.02-µm Anodisc filters (Whatman, UK), and stained with 10× SYBR Gold (Invitrogen, USA). VLPs were then counted using an epifluorescence microscope (BX51-FL, Olympus, Japan). The phage concentrate was stored in a 40% glycerol solution at −70 °C. The stock solution was 10-fold diluted and used as an internal standard.

### VLP enrichment, DNA extraction, and shotgun metagenome sequencing

Samples were homogenized with 100–300 ml of SM buffer in a sterile plastic bag (BagFilter^®^, Interscience, France) using a stomacher at speed 4 (BagMixer^®^ 400 CC, Interscience, France) for 2 min. The homogenate was centrifuged at 8,000 ×g for 10 min at 4 °C, and the supernatant was transferred into a new tube. The pellet was stored at −70 °C for bacterial metagenome DNA extraction. This step was repeated once or twice to ensure complete collection of VLPs and microbial cells. A diluted solution of *E. coli* T4 phage was added to the supernatant in total amounts ranging from 5.0 × 10^5^ to 1.0 × 10^8^ VLPs (Supplementary Table 1). The supernatant was sequentially filtered through 0.45-µm and 0.2-µm CA syringe filters (Sartorius, Germany). The filtrate was mixed with 10% (wt/vol) polyethylene glycol 8000 (Sigma-Aldrich, USA) and 1 M NaCl (Bioneer, South Korea), and incubated at 4 °C for 15 hr. The mixture was centrifuged at 13,000 ×g for 30 min at 4 °C, and the resulting pellet was resuspended in 5 ml SM buffer. The resuspension was further concentrated using 100 kDa MWCO centrifugal filters (Amicon^Ⓡ^ Ultra-15 Sigma-Aldrich, USA). The concentrate was treated with 20 µl lysozyme (10 mg/ml, Biosolution, South Korea), and incubated at 37 °C for 30 min. It was further incubated with 0.2 volume of chloroform at 4 °C for 30 min, with vortexing every 5 min. Chloroform was removed by centrifugation at 4,500 ×g for 10 min at 4 °C. The concentrate was treated with three units of DNase Ⅰ (Sigma-Aldrich, USA) and RNase A (Bioneer, South Korea) and incubated at 37 °C for up to 3 hr to remove free nucleic acids. A stop solution was then added and incubated at 65 °C for 15 min. Total VLP DNA was extracted by treating 30 µl of 10% SDS (Sigma-Aldrich, USA) and 3 µl protease K (20 mg/ml, 600 mAU/ml, Bioneer, South Korea) at 55 °C for 1 hr. Subsequently, 100 µl of 5 M NaCl and 10% (v/v) cetyltrimethylammonium bromide buffer were added and incubated at 65 °C for 10 min. An equal volume of phenol/chloroform/isopropanol (25:24:1, Sigma-Aldrich, USA) was added, and the aqueous solution was collected by centrifugation at 8,000 ×g for 10 min at room temperature. Chloroform was repeated in the same manner. The supernatant was then incubated with 0.1 volume of 3 M sodium acetate (Bioneer, South Korea), 1 µl glycogen (20 mg/ml, Sigma-Aldrich, USA), and 0.7 volume of isopropanol (Sigma-Aldrich, USA) at −20 °C for 2 hr, followed by centrifugation at 13,000 ×g for 20 min at 4 °C. The pellet was washed with 70% cold ethanol and dried for 20 min at room temperature. Total VLP DNA was eluted in 200 µl RNase-free dH_2_O (Bioneer, South Korea) by incubating at 55 °C for 5 min. The VLP DNA was further purified using QIAamp MinElute Virus Spin Kit (Qiagen, Germany). Negative controls were prepared to detect possible contamination from the reagents in the commercial kits, but no DNA was detected. Sequencing libraries were constructed using the Accel-NGS 1S Plus DNA Library kit (Swift Biosciences, USA), according to the manufacturer’s instruction. Shotgun metagenome sequencing was performed on an Illumina NovaSeq 6000 sequencer (2 × 150 bp) (Illumina, USA).

### Bacterial DNA extraction and amplicon sequencing

The pellet obtained from the sample homogenate was resuspended in 5 ml SM buffer, and were filtered through 5.0-µm syringe filters (Sartorius, Germany). The filtrate was centrifuged at 8,000 ×g for 10 min at 4 °C, and bacterial DNA was extracted from the pellet using QIAamp^®^ Fast DNA Stool Mini Kit (Qiagen, Germany) with repeated bead beating at 5.5 m/s for 30 sec in a FastPrep^®^-24 instrument (MP biomedicals, USA). Negative controls were also included to detect possible contamination from the reagents in commercial kits. The V5-V6 regions of bacterial 16S rRNA gene were amplified with chloroplast-excluding primers, combined with Illumina sequencing overhang adapters using AccuPower PCR Premix (Bioneer, South Korea), as described previously^9^. Amplicons were prepared in triplicates and purified using the QIAquick PCR purification kit (Qiagen, USA). The 16S amplicons were sequenced on an Illumina MiSeq sequencer (2 × 300 bp) (Illumina, USA).

### Preprocessing of VLP metagenome data

Adapter sequences and low-quality reads were removed from raw reads using Fastp v0.23.1^83^ with default settings. Twenty base pairs were trimmed from the end of R1 reads and the start of R2 reads, according to the manufacturer’s recommendation (Swift Biosciences, USA). Duplicates were removed using Fastp v0.23.1 with the setting “*--dup_calc_accuracy 6*”. Reads considered being derived from plants were removed by mapping to representative genomes of fresh vegetables (broccoli, GCF_000695525.1; perilla, GCA_004303085.1; spinach, GCF_002007265.1; lettuce, GCF_002870075.1) using Bowtie2 with *--local* mode. Reads that were taxonomically assigned into the domains *Metazoan* and *Viridiplantae* using Kraken2^84^ were further removed. High-quality reads were aligned to the T4 phage genome (HM137666) using Bowtie2 with default settings. The aligned reads and the remaining reads were separated and assembled individually for each sample using metaSPAdes v3.13.1^85^.

### Viral operational taxonomic unit

Large contigs (≥2.5 kb) were retained from the assemblies. These contigs were classified as viral, if they met any of the following criteria: (1) classified into categories 1, 2, 4, or 5 by VirSorter v1.0.6^86^; (2) identified as viruses by VirFinder v1.2.1^87^ with a score ≥ 0.9 and *p* < 0.05; (3) classified into categories 3 or 6 by VirSorter and identified as viruses by VirFinder with a score ≥ 0.7 and *p* < 0.05; (4) classified into categories 3 or 6 by VirSorter and identified as viruses by VIBRANT v1.2.1; (5) identified as viruses by both VirFinder with a score ≥ 0.7 and *p* < 0.05, and VIBRANT; or (6) classified as viruses or unclassified with less than 40% open reading frames assigned to Bacteria, Archaea, or Eukaryotes using CAT v4.8.1^88^. VirSorter and VIBRANT were executed in *–virome* mode. Circular contigs, where the start and end of the contig were identical, were also classified as viral. Exact duplicates were removed using CD-HIT-EST v4.8.1. The completeness of non-redundant viral contigs (complete, high-, medium- and low-quality) were assessed using CheckV v0.7.0. Viral contigs were then clustered into vOTUs using the ANIm of dRep v3.0.1^89^, if they shared ≥95% average nucleotide identity over 85% of the alingment fraction. Representative genomes for the vOTUs were selected based on completeness and centrality. High-quality reads were mapped to the representative genomes using BBMap v38.70 with default parameters, and the genomes with ≥75% breadth of coverage and ≥1× depth of coverage were retained. The number of mapped reads was normalized using the FPKM method, with normalized values rounded down. Alpha-diversity was estimated using the number of vOTUs and Shannon index. Beta-diversity was measured based on Bray-Curtis dissimilarity with log-transformation.

### Viral enrichment estimation

Viral enrichment was assessed using ViromeQC with ‘*-w* environmental’ option. The viral enrichment score was calculated at each stage of sequence processing (raw reads, preprocessed reads, high-quality reads, reads mapped to large contigs, and reads mapped to viral contigs) by counting the number of reads corresponding to large and small subunit ribosomal RNA gene, as well as single-copy universal markers of bacterial housekeeping genes.

### Bacterial 16S rRNA gene analysis

16S rRNA gene amplicon sequences were analyzed using QIIME2 v2021.08. Briefly, Illumina adapter sequences were trimmed, and ASVs were generated using DADA2^90^. DADA2 corrected low-quality bases and errors, merged paired-end reads, and removed chimeric sequences. Taxonomic identification of the ASVs was performed using Naïve Bayes classifier trained on non-redundant (99% identity) SILVA small subunit rRNA database (version 138). ASVs assigned to eukaryotes, archaea, mitochondria, chloroplast, or unclassified were excluded. Sequencing depth was evenly rarefied to 20,000 sequences, and ASVs with fewer than 10 sequences were further removed. Alpha- and beta-diversity were analyzed in the same manner as the vOTU analysis.

### Quantification of total viral and bacterial abundances

The number of total viral genomes was calculated from the vOTU table using FPKM values, with the following equation: Total number of viral genomes = (Number of T4 phage VLPs × ∑ FPKM values of viral contigs) / ∑ FPKM values of T4 phage contigs. The total number of bacterial genomes was estimated by quantifying 16S rRNA gene copies using quantitative PCR with chloroplast-excluding primers^91^. The 16S rRNA gene sequences of *E*. *coli* K12 were used as a standard. The number of 16S rRNA gene copies was then normalized based on the average number of 16S rRNA gene copies per genome in *Gammaproteobacteria*^92^ whose ASVs were the most abundant (51.6 ± 28.2% on average) in the bacterial microbiome of fresh vegetables.

### Viral clusters and taxonomic identification

The viral genomes from NCBI RefSeq Viral Genome (2022-03-10 release) and IMG/VR (2020-10-12 release) were selected by comparing them with the representative sequences of the vOTUs (BLASTn, *E*-value < 10^-5^). A gene-sharing network was constructed with all vOTU genomes using vConTACT2, with an inflation of 1.5 for protein clusters and 1 for VCs. The abundance of VCs was determined by summing the abundances of all vOTUs within each VC. The genus-level taxonomy of VCs (≥ two members) was determined based on the taxonomy of viral genomes from NCBI RefSeq Viral Genome and IMG/VR. Specifically, the protein sequences of the vOTUs were compared with those from NCBI RefSeq Viral Genome and IMG/VR within each VC using BLASTp (*E*-value < 10^-5^, bit-score > 50). The best BLAST results were then used to identify the genus-level taxonomy through the lowest common ancestor algorithm in MEGAN6. vOTUs categorized as “Clustered” were used to determine genus-level taxonomy. vOTUs categorized as “Clustered”, “Clustered/Singleton”, or “Overlap” were considered for family-level taxonomy. The majority rule was employed to determine the family-level taxonomy of vOTUs assigned to the “Overlap” category comprising multiple VCs. The family-level taxonomy of VCs categorized as “Clustered/Singleton” followed the taxonomy of their pre-VCs. The taxonomy of vOTUs were described according to the ICTV taxonomy (2019 release)^32^.

### Viral lifestyle prediction

The infection lifestyle was predicted using BACPHLIP^93^ and CheckV. Temperate phages were identified by detecting lysogeny-associated genes, such as integrase and recombinase, using BACPHLIP, or by finding integrated viral sequences using CheckV. vOTUs that did not show lysogenic signals were classified as virulent phages.

### Prokaryotic host prediction

First, bacterial hosts were predicted using iPHoP v1.3.2 which employs random forest classifier trained on CRISPR spacer identification, *k*-mer similarity, and homologous sequence match with provided Sept_2021_pub database based on positive predictive value (designated as confidence score ≥ 90). When multiple hosts were predicted, one with the highest score was selected. Subsequently, CRISPR spacer identification and tRNA sequence similarity were employed to identify bacterial hosts for vOTUs when host predictions from iPHoP were not available. CRISPR spacer sequences from Dion *et al*.^94^ were used to query vOTUs using BLASTn (≥95% identity, ≥95% coverage, and ≤2 mismatches). Additional CRISPR spacer sequences were obtained by extracting them from the bacterial genomes of GTDB (release 214)^95^ and GEM databases^96^ using MinCED v0.4.2^97^ with default parameters, and used in the same manner. tRNA sequences were identified from the genoems of vOTUs using tRNAscan-SE v1.4 with default parameters. The tRNA sequences were compared with bacterial genomes from the GTDB and GEM databases using BLASTn (≥100% identity, ≥100% coverage). Hosts were assigned to vOTUs only if the taxonomy of all hits was consistent. Hosts were not assigned if they were not observed in the 16S rRNA gene data.

### Auxiliary metabolic gene annotation

AMGs were identified using DRAM-v and VIBRANT. For DRAM-v annotation, VirSorter2 was run on all vOTUs with ‘*--prep-for-dramv*’ option to identify viral genes and viral hallmark genes. The results were then used for DRAM-v analysis. Putative AMGs were selected with high auxiliary scores (category 1-3). For VIBRANT annotation, AMGs were identified from vOTUs with a high enrichment of viral genes. Only putative AMGs categorized as metabolic pathway and sulfur relay system in KEGG annotations were considered. Based on the report of Luo *et al*.^20^, AMGs related to ribosomal protein, glycosyl transferase, glycoside hydrolase, methyltransferase, peptidase, nucleotide and amino acid metabolism were further excluded, as these genes are essential for viral functioning and are likely to be illegitimate AMGs.

### Comparison with public gut virome databases and human gut VLP metagenome datasets

Raw reads of human gut VLP metagenome datasets (13 studies, n = 1,306 subjects) were collected from NCBI Sequence Read Archive (Supplementary Table 5). Illumina adapter sequences and low-quality reads were trimmed using Trimmomatic v0.39 (ILLUMINACLIP:TruSeq3-PE-2.fa:2:30:10, LEADING:20, TRAILING:20, SLIDINGWINDOW:4:20, MINLEN:100-240). Exact duplicates were removed using PRINSEQ v0.20.4. Reads mapped to NCBI RefSeq human genome (GCF_000001405) were removed using Bowtie2. High-quality reads were then mapped to the genome sequences of vOTUs and crAssphage (JQ995537) using BBMap v38.70 with settings of 95% identity and ‘*ambiguous=all*’. vOTUs with fewer than 10 paired-end reads, or with a depth of coverage of less than 0.5× across all samples considered potential false positives. The number of mapped reads per vOTU was normalized using the FPKM method, accounting for the read length of each study. Alpha- and beta-diversity were analyzed in the same manner as the vOTU analysis. vOTUs that were significantly enriched or depleted in either healthy or diseased group were identified using a generalized linear-mixed effects model in MaAsLin2^98^. Viral genomes from the GVD, GPD, MGV, and IMG/VR databases were compared at the genus-level with those of vOTUs using vConTACT2. Species-level comparisons between viral genomes from these databases and vOTUs were determined by calculating average nucleotide values using “anicalc.py” in CheckV.

### Associations between dietary patterns and gut-associated vegetable vOTUs

We recruited a total of 53 healthy individuals (Supplementary Table 7), their dietary intake patterns were evaluated using both short-term and long-term methods (The Institutional Review Board of Chungnam National University, 202012-SB-169-01). For short-term assessment, participants reported all foods and beverages consumed over a period of three days. Fecal samples were collected on the subsequent day following the completion of the final meal record, and their fecal samples were processed in the same manner as the experimental procedure of the vegetable virome. The meal records were categorized into 16 food groups by modifying the classification system provided by the Rural Development Administration in Korea, and then converted to serving sizes in accordance with the established protocols^99^. For long-term assessment, participants completed a food frequency questionnaire comprising 127 food items, designed specifically for Korean adult^100^. The questionnaire was evaluated the frequency and serving size of each item consumed over the past 12 months, subsequently classifying the 127 food items into 20 food groups. The intake amounts of the food groups were transformed into centered log ratios. Correlations between food groups and gut-associated vOTUs were determined using PERMANOVA and db-RDA analysis. The intake values of four short-term food groups (vegetables, fruits, potatoes, and legumes) were further categorized into three groups (high, medium, and low), whose sample number was evenly distributed across the groups. Gut-associated vOTUs that were enriched or depleted in each group were identified using MaAsLin2, with adjustments for fruit, potato, and legume.

### Statistical analysis

*P*-value of < 0.05 was considered significant. One-tailed and two-tailed Mann-Whitney U tests, one-way ANOVA with Tukey’s post-hoc test, and Kruskal-Wallis test with Dunn’s post-hoc test were performed using GraphPad Prime v5.01 for Windows (GraphPad Software, USA). PCoA was performed based on Jaccard distance and Bray-Curtis dissimilarity using R package ‘vegan’. Statistical significance for distance matrice was assessed using the ‘adonis’ function of ‘vegan’ package, with 999 permutations. The best-fit models constrained to host and environmental factors were selected using ordiR2step with forward selection in R package ‘vegan’, with 999 permutations. The percentages of explained variances for host and environmental factors were calculated using db-RDA of R package ‘vegan’. vOTUs differentially abundant among groups were determined using MaAsLin2 with a prevalence of ≥ 10%.

### Data availability

The EMBL-EBI accession number for the sequences of the virome and 16S rRNA gene data is PRJEB79316.

## Acknowledgments

This research was supported by the National Research Foundation of Korea (NRF) grants funded by the Ministry of Education (2018R1D1A3B07050366) and the Ministry of Science and ICT (MSIT) (No.2019R1C1C1009664, No.2021R1F1A1064222, and No.2020R1A5A8017671).

## Author contributions

J.W.P., E.J.P. and M.S.K. conceived and designed the experiments; J.W.P., Y.E.Y., and S.A.I. performed the experiments; J.W.P., Y.E.Y. and M.S.K. analyzed the data; J.A.C, S.I.Y., E.J.P., and M.S.K. contributed materials/analysis tools; J.W.P., E.J.P. and M.S.K. wrote the manuscript.

## Supplementary Information

**Extended Data Fig. 1.**
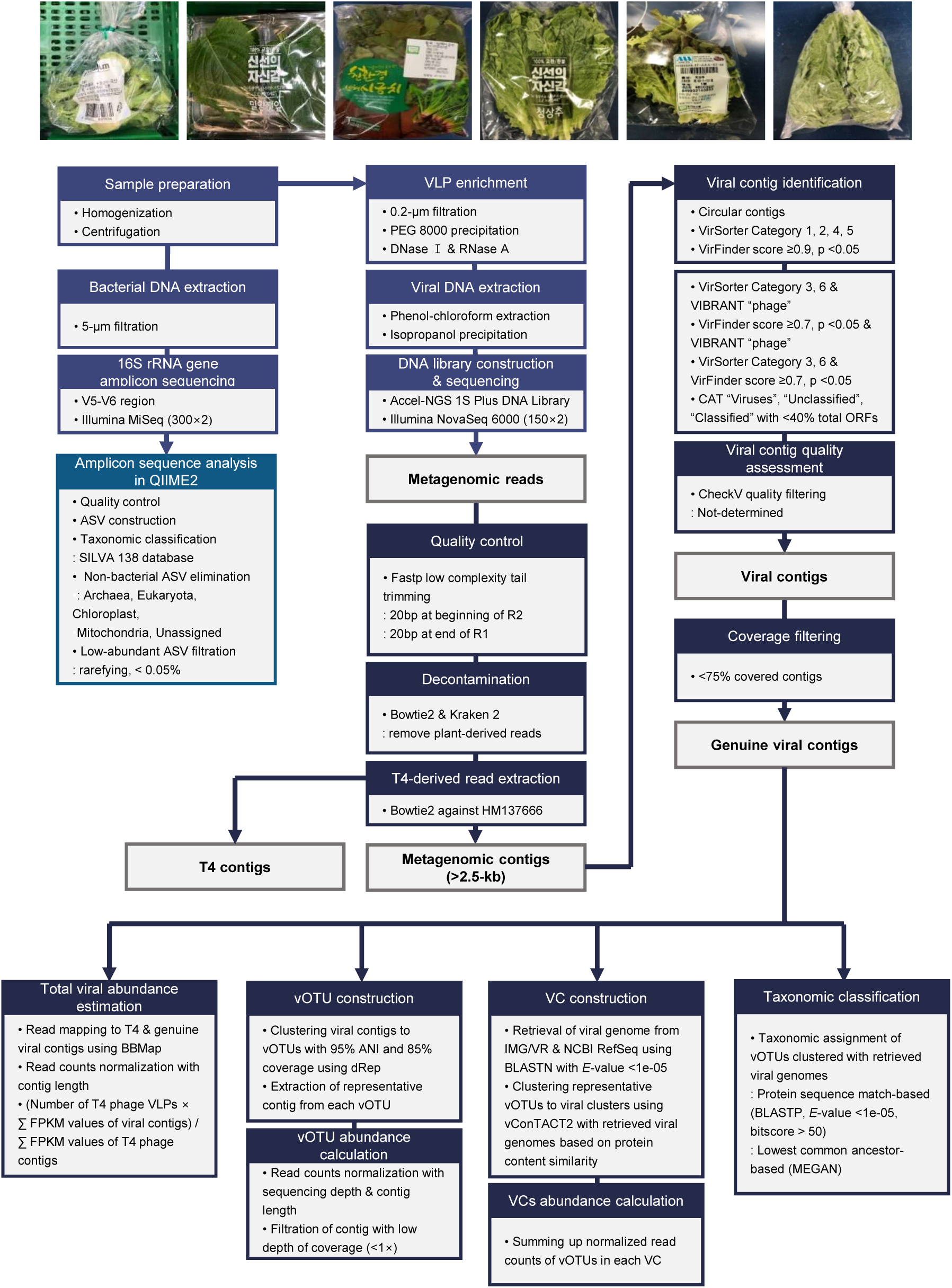
Analytical workflow for viral and bacterial metagenome sequences.

**Extended Data Fig. 2.**
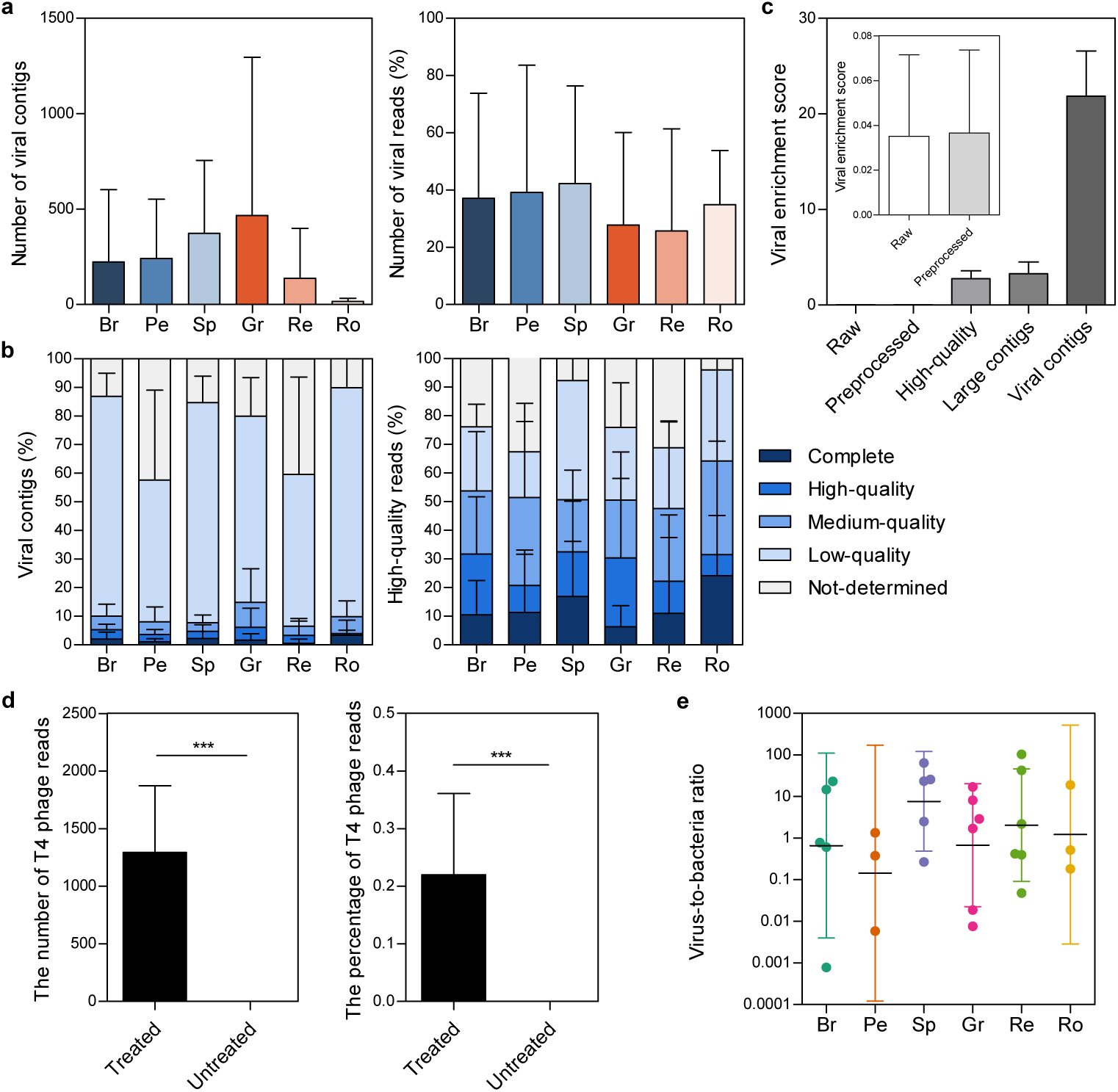
Reconstruction of viral genomes from fresh vegetable VLP metagenomes. **a,** The number (left panel) and percentage (right panel) of viral contigs obtained from the VLP metagenomes. **b,** The percentages of viral contigs classified as complete, high-, medium-, low-quality, and not-determined (left panel), and their abundance (right panel). **c,** Viral enrichment scores for each sequence processing stage. **d,** The number and percentage of reads mapped to *E. coli* T4 phage genome. **e,** Virus-to-bacteria ratio across the vegetable types are shown. Data are presented as geometric mean ± geometric S.D. Statistical significance was determined by two-tailed Mann-Whitney test. Symbol: ***, *P* < 0.001.

**Extended Data Fig. 3.**
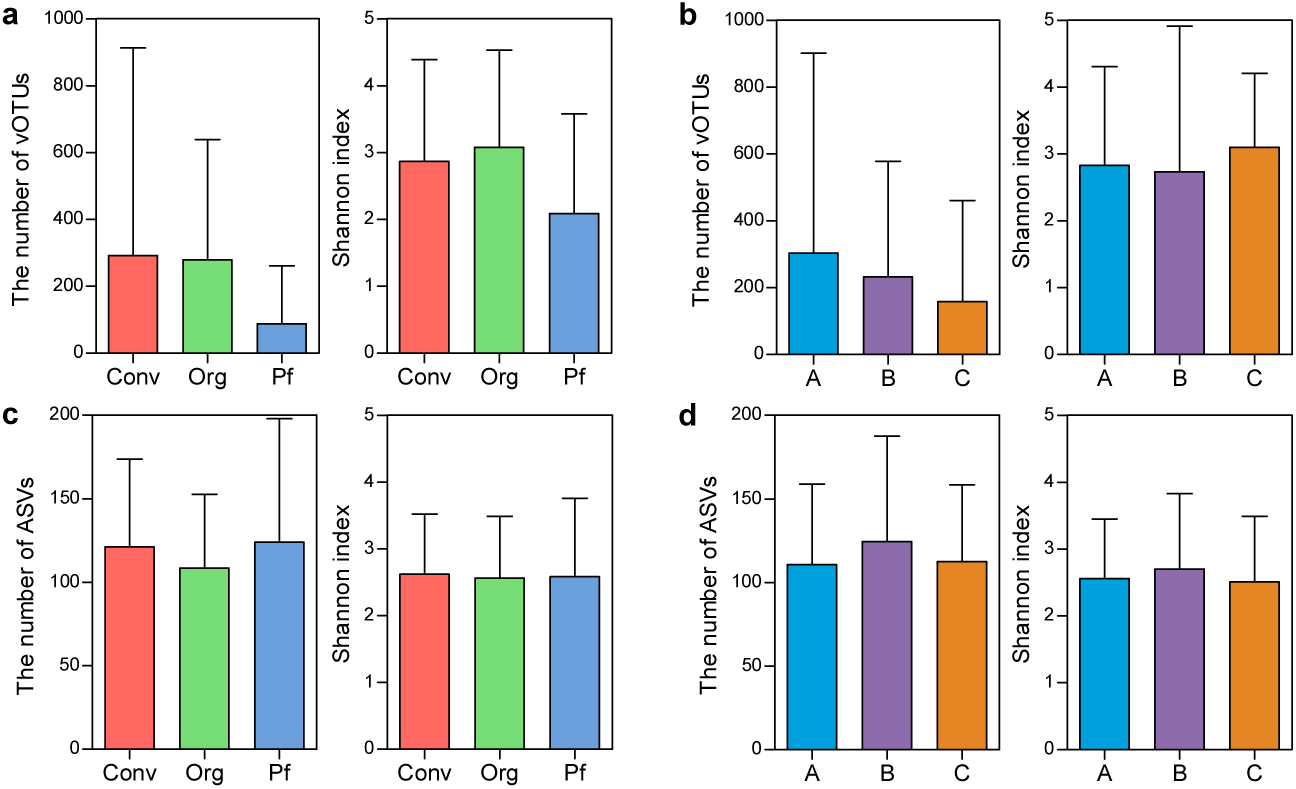
Species-level diversity of vegetable viral and bacterial microbiomes, related to Fig. 2. **a,b,** The number of observed vOTUs (left panel) and Shannon index (right panel) among the farming methods **(a)** and vendors **(b)**. **c,d,** The number of observed ASVs (left panel) and Shannon index (right panel) among the farming methods **(c)** and vendors **(d)**. All data are presented as mean ± S.D.

**Extended Data Fig. 4.**
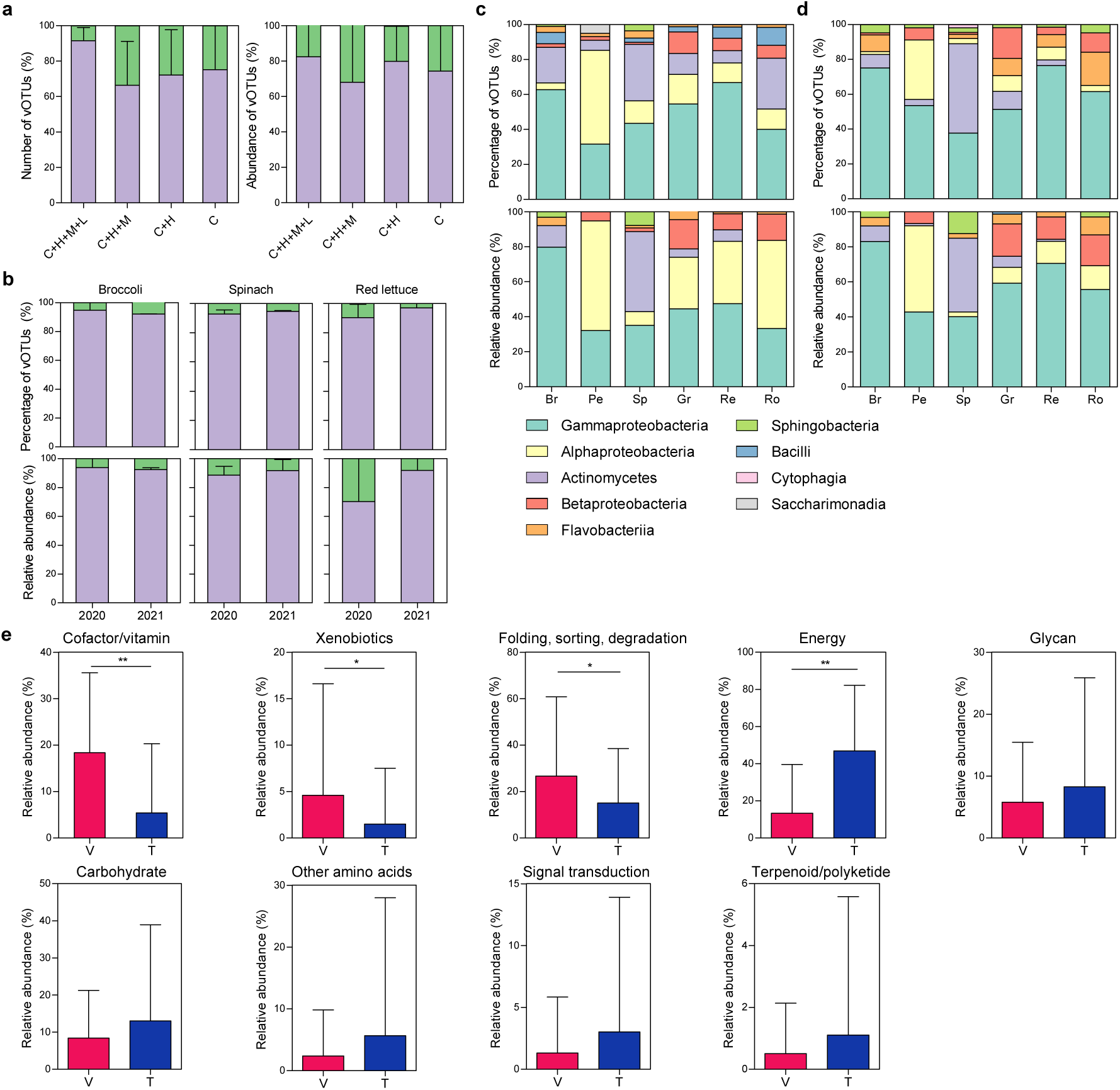
Distribution of viral lifestyles and AMGs. **a,** The percentage of virulent and temperate vOTUs (left panel), and their relative abundance (right panel) according to different genome qualities (C, complete-quality; H, high-quality; M, medium-quality; and L, low-quality). **b,** The percentage of virulent and temperate vOTUs (upper panel), and their relative abundance (lower panel) among the sampling periods. **c,d,** The percentage of bacterial hosts (upper panel) and their relative abundance (lower panel) of virulent **(c)** and temperate vOTUs **(d)** across the vegetable types. Only bacterial classes with an average percentage ≥ 1% are shown. **e,** The relative abundance of AMG functions was compared between virulent and temperate vOTUs. Data are presented as mean ± S.D. Statistical significance was determined by the Mann-Whitney test. Symbol: *, *P* < 0.05; and **, *P* < 0.01.

**Extended Data Fig. 5.**
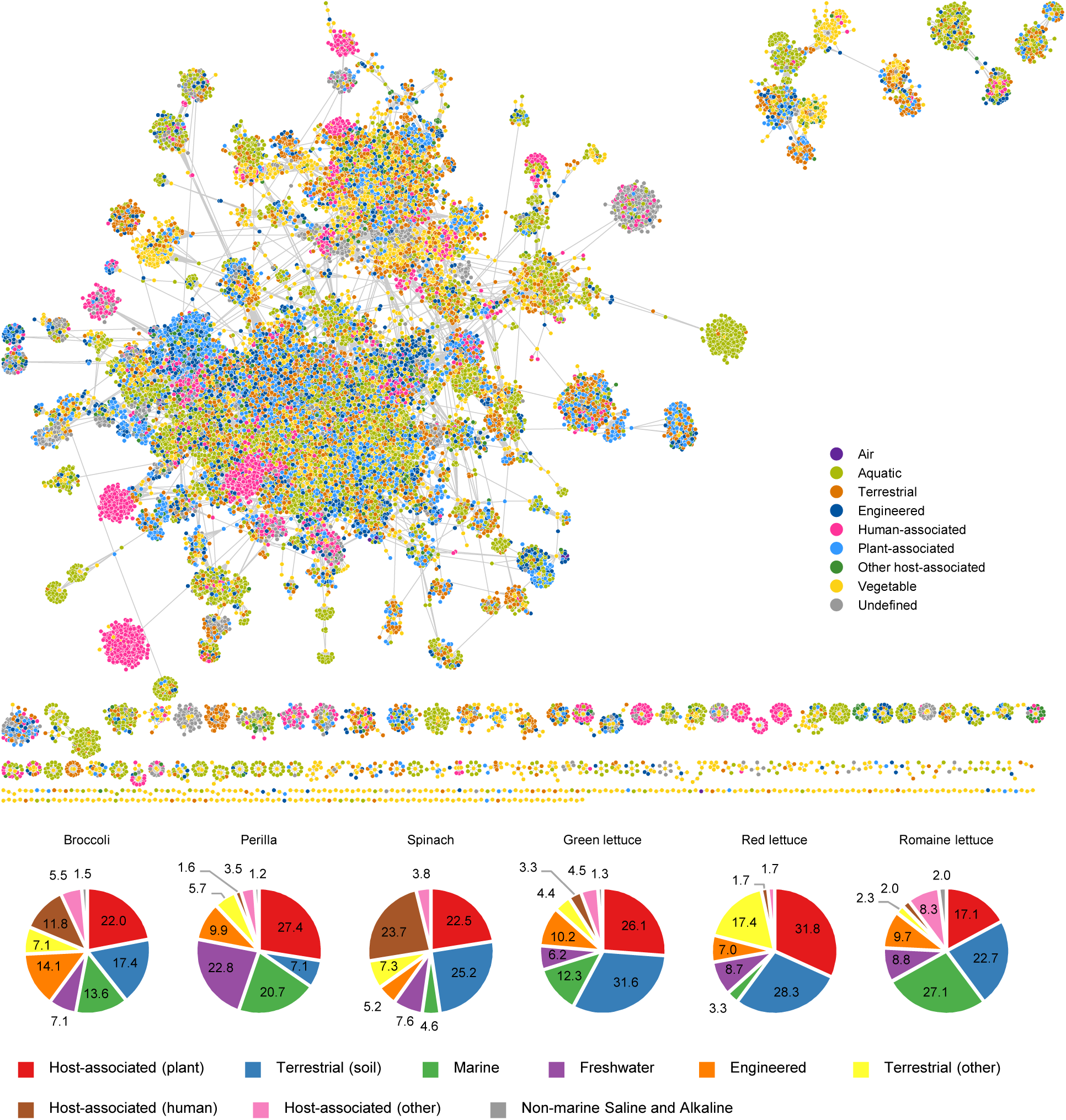
Genus-level associations between viral genomes from fresh vegetables and various ecosystems. A gene-sharing network was constructed with the genomes of vegetable vOTUs and IMG/VR. Pie charts show the percentage of environmental associations between the vegetable virome and nine ecosystems. Only associations with vOTUs ≥ 1% are shown.

**Extended Data Fig. 6.**
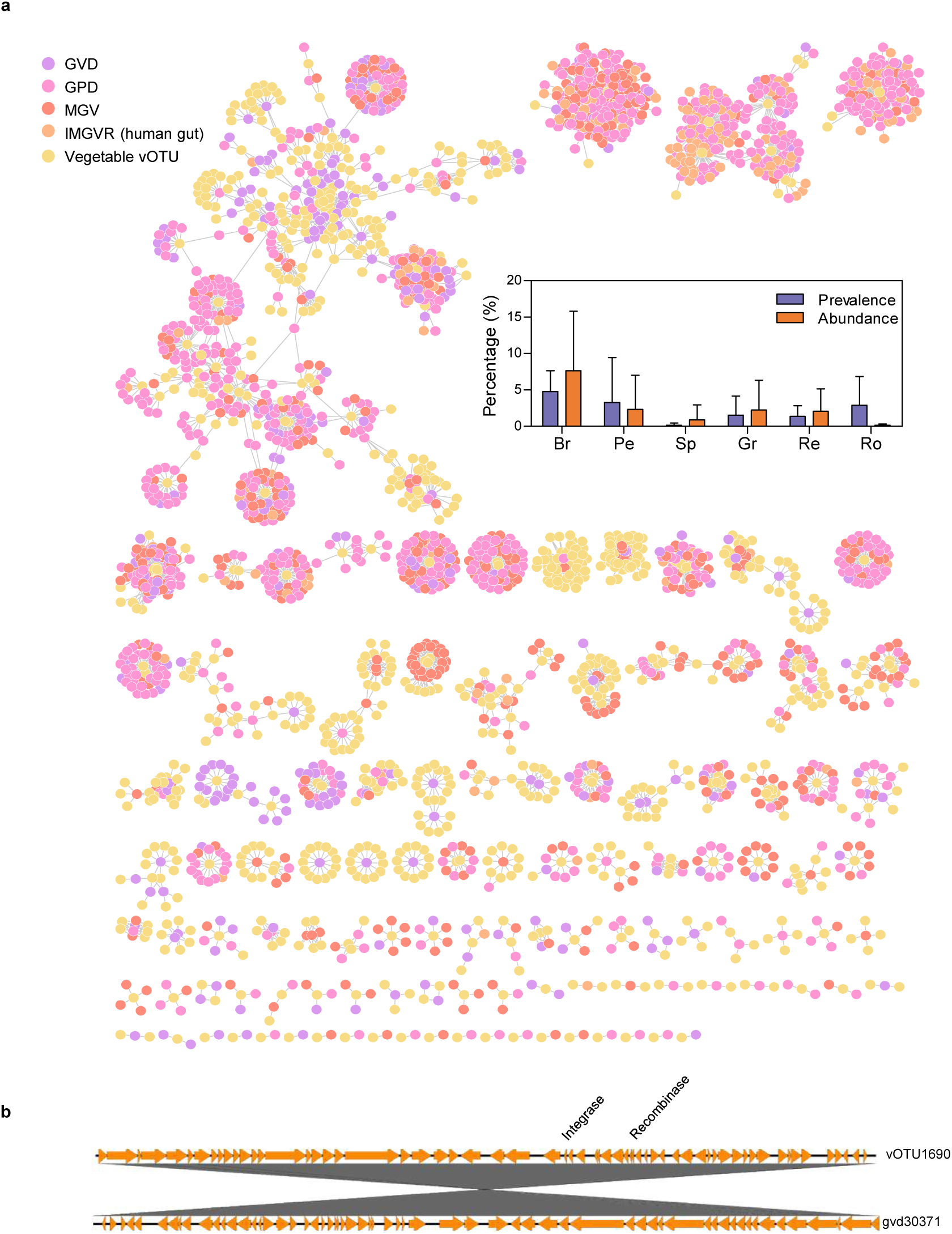
Genus-level associations between the viromes from fresh vegetables and four gut virome databases. **a,** A gene-sharing network was constructed with the genomes of vegetable vOTUs, IMG/VR, and human gut virome databases (GVD, GPD, and MGV). Box plot shows the prevalence and percentage of gut-associated vOTUs at the genus level. **b,** Genomic comparison between the genomes of gut-associated vOTU1690 (upper) and GVD30371 (lower).

**Extended Data Fig. 7.**
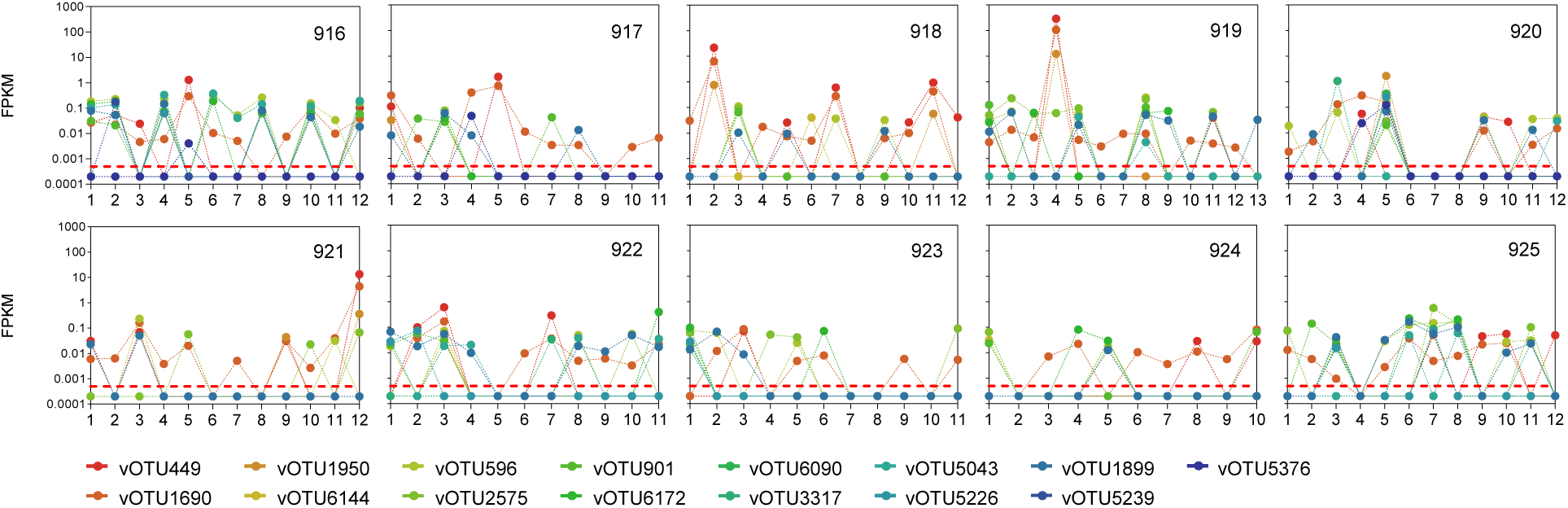
Dynamics of the abundance of 15 global vOTUs in the longitudinal samples of Shkoporov *et al*. (2019)

**Extended Data Fig. 8.**
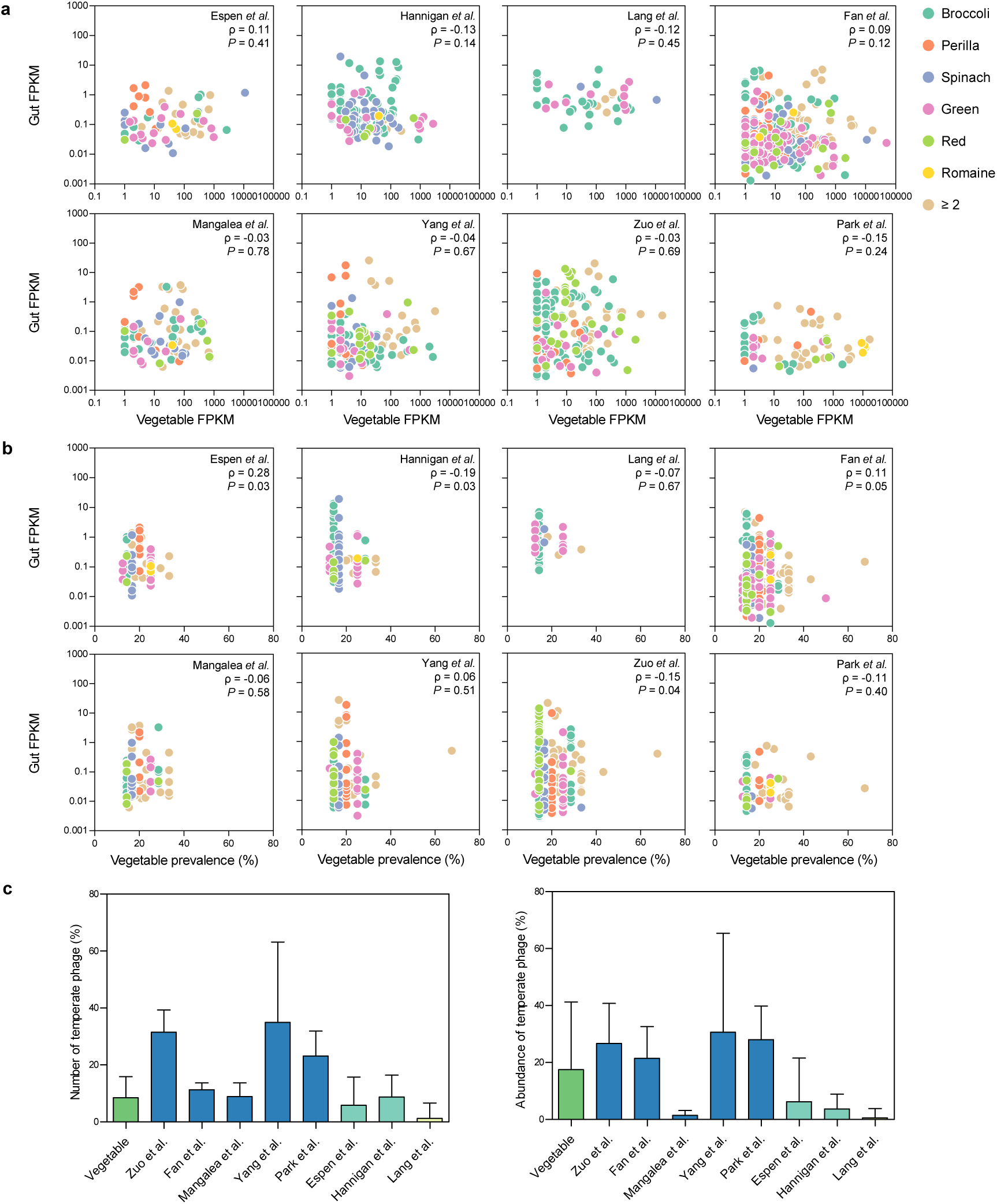
Selective transmission of vegetable viruses to the human gut. **a,** Spearman correlations between the abundance of gut-associated vOTUs in vegetable samples and in the VLP metagenomes. **b,** Spearman correlations between the prevalence of gut-associated vOTUs in vegetable samples and the abundance of gut-associated vOTUs in the gut VLP metagenomes. **c,** The percentage (left panel) and relative abundance (right panel) of gut-associated virulent and temperate vOTUs.

**Supplementary Table S1** Sample preparation information of fresh vegetable produces in present study.

**Supplementary Table S2** Completeness, taxonomic, lifestyle, and host bacteria information of 6,363 vOTUs.

**Supplementary Table S3** Annotation of viral AMGs.

**Supplementary Table S4** Genomic comparison between vegetable vOTUs and GVDs. Genomic similarity expressed as average nucleotide identity.

**Supplementary Table S5** Detailed information about the 13 huVLPs.

**Supplementary Table S6** Distribution of gut-associated vOTUs in the huVLPs.

**Supplementary Table S7** Detailed information about the 53 subjects.

## References

1 Carter, M. M. et al. Ultra-deep sequencing of Hadza hunter-gatherers recovers vanishing gut microbes. Cell 186, 3111–3124 (2023).

2 Moraïs, S. et al. Cryptic diversity of cellulose-degrading gut bacteria in industrialized humans. Science 383, eadj9223 (2024).

3 Rothschild, D. et al. Environment dominates over host genetics in shaping human gut microbiota. Nature 555, 210–215 (2018).

4 Valles-Colomer, M. et al. The person-to-person transmission landscape of the gut and oral microbiomes. Nature 614, 125–135 (2023).

5 Desai, M. S. et al. A dietary fiber-deprived gut microbiota degrades the colonic mucus barrier and enhances pathogen susceptibility. Cell 167, 1339–1353 (2016).

6 Sonnenburg, E. D. et al. Diet-induced extinctions in the gut microbiota compound over generations. Nature 529, 212–215 (2016).

7 Leff, J. W. & Fierer, N. Bacterial communities associated with the surfaces of fresh fruits and vegetables. PloS One 8, e59310 (2013).

8 Kim, M.-S. & Park, E.-J. Composition and variability of core phyllosphere fungal mycobiota on field-grown broccoli. *Environ*. Microbiome 18, 15 (2023).

9 Kim, M.-S., Bae, J.-W. & Park, E.-J. Geographic and host-associated variations in bacterial communities on the floret surfaces of field-grown broccoli. Appl. Environ. Microbiol. 84, e02837–02817 (2018).

10 Dees, M. W., Lysøe, E., Nordskog, B. & Brurberg, M. B. Bacterial communities associated with surfaces of leafy greens: shift in composition and decrease in richness over time. Appl. Environ. Microbiol. 81, 1530–1539 (2015).

11 Kim, M.-S. & Park, E.-J. Postharvest-induced microbiota remodeling increases fungal diversity in the phyllosphere mycobiota of broccoli florets. Postharvest Biol. Technol. 181, 111693 (2021).

12 Kim, M. S., Bae, J. W. & Park, E. J. Postharvest processing decreases the richness of bacterial taxa in the phyllosphere of broccoli. J. Appl. Microbiol. 125, 295–305 (2018).

13 Berg, G. et al. Plant microbial diversity is suggested as the key to future biocontrol and health trends. FEMS Microbiol. Ecol. 93, fix050 (2017).

14 Kusstatscher, P. et al. Microbiome approaches provide the key to biologically control postharvest pathogens and storability of fruits and vegetables. FEMS Microbiol. Ecol. 96, fiaa119 (2020).

15 Milani, C. et al. Colonization of the human gut by bovine bacteria present in Parmesan cheese. Nat. Commun. 10, 1286 (2019).

16 Pasolli, E. et al. Large-scale genome-wide analysis links lactic acid bacteria from food with the gut microbiome. Nat. Commun. 11, 2610 (2020).

17 Wicaksono, W. A. et al. The edible plant microbiome: evidence for the occurrence of fruit and vegetable bacteria in the human gut. Gut Microbes 15, 2258565 (2023).

18 Cobián Güemes, A. G., et al. Viruses as winners in the game of life. Annu. Rev. Virol. 3, 197–214 (2016).

19 Hsu, B. B. et al. Dynamic modulation of the gut microbiota and metabolome by bacteriophages in a mouse model. Cell Host Microbe 25, 803–814. (2019).

20 Luo, X.-Q. et al. Viral community-wide auxiliary metabolic genes differ by lifestyles, habitats, and hosts. Microbiome 10, 190 (2022).

21 Sinha, A. et al. Transplantation of bacteriophages from ulcerative colitis patients shifts the gut bacteriome and exacerbates the severity of DSS colitis. Microbiome 10, 105 (2022).

22 Ritz, N. L. et al. The gut virome is associated with stress-induced changes in behaviour and immune responses in mice. Nat. Microbiol. 9, 359–376 (2024).

23 Roux, S. et al. Towards quantitative viromics for both double-stranded and single-stranded DNA viruses. PeerJ 4, e2777 (2016).

24 Norman, J. M. et al. Disease-specific alterations in the enteric virome in inflammatory bowel disease. Cell 160, 447–460 (2015).

25 Kim, K.-H. & Bae, J.-W. Amplification methods bias metagenomic libraries of uncultured single-stranded and double-stranded DNA viruses. Appl. Environ. Microbiol. 77, 7663–7668 (2011).

26 Nayfach, S. et al. CheckV assesses the quality and completeness of metagenome-assembled viral genomes. Nat. Biotechnol. 39, 578–585 (2021).

27 Zolfo, M. et al. Detecting contamination in viromes using ViromeQC. Nat. Biotechnol. 37, 1408–1412 (2019).

28 López-García, P. et al. Metagenome-derived virus-microbe ratios across ecosystems. ISME J. 17, 1552–1563 (2023).

29 Knowles, B. et al. Lytic to temperate switching of viral communities. Nature 531, 466–470 (2016).

30 Lindow, S. E. & Brandl, M. T. Microbiology of the phyllosphere. Appl. Environ. Microbiol. 69, 1875–1883 (2003).

31 Roux, S. et al. Minimum information about an uncultivated virus genome (MIUViG). Nat. Biotechnol. 37, 29–37 (2019).

32 Adriaenssens, E. M. Phage diversity in the human gut microbiome: A taxonomist’s perspective. mSystems 6, e00799–21 (2021).

33 Bin Jang, H., et al. Taxonomic assignment of uncultivated prokaryotic virus genomes is enabled by gene-sharing networks. Nat. Biotechnol. 37, 632–639 (2019).

34 Roux, S. et al. iPHoP: An integrated machine learning framework to maximize host prediction for metagenome-derived viruses of archaea and bacteria. PLoS. Biol. 21, e3002083 (2023).

35 Shaffer, M. et al. DRAM for distilling microbial metabolism to automate the curation of microbiome function. Nucleic Acids Res. 48, 8883–8900 (2020).

36 Kieft, K., Zhou, Z. & Anantharaman, K. VIBRANT: automated recovery, annotation and curation of microbial viruses, and evaluation of viral community function from genomic sequences. Microbiome 8, 1–23 (2020).

37 Yuan, Y. et al. Discovery of novel bacterial queuine salvage enzymes and pathways in human pathogens. Proc. Natl. Acad. Sci. U. S. A. 116, 19126–19135 (2019).

38 Kredich, N. M. Biosynthesis of cysteine. EcoSal Plus 3, 10.1128/ecosalplus.3.6.1.11 (2008).

39 Zhang, T. et al. RNA viral community in human feces: prevalence of plant pathogenic viruses. PLoS. Biol. 4, e3 (2006).

40 David, L. A. et al. Diet rapidly and reproducibly alters the human gut microbiome. Nature 505, 559–563 (2014).

41 Van Espen, L., et al. A previously undescribed highly prevalent phage identified in a Danish Enteric Virome Catalog. mSystems 6, 10.21203/rs.3.rs-273865/v1 (2021).

42 Camarillo-Guerrero, L. F., Almeida, A., Rangel-Pineros, G., Finn, R. D. & Lawley, T. D. Massive expansion of human gut bacteriophage diversity. Cell 184, 1098–1109 (2021).

43 Gregory, A. C. et al. The gut virome database reveals age-dependent patterns of virome diversity in the human gut. Cell Host Microbe 28, 724–740 (2020).

44 Nayfach, S. et al. Metagenomic compendium of 189,680 DNA viruses from the human gut microbiome. Nat. Microbiol. 6, 960–970 (2021).

45 Li, J., Yang, F., Xiao, M. & Li, A. Advances and challenges in cataloging the human gut virome. Cell Host Microbe 30, 908–916 (2022).

46 Pinto, Y., Chakraborty, M., Jain, N. & Bhatt, A. S. Phage-inclusive profiling of human gut microbiomes with Phanta. Nat. Biotechnol. 42, 651–662 (2024).

47 Wan, Y. et al. Alterations in fecal virome and bacteriome virome interplay in children with autism spectrum disorder. Cell Rep. Med. 5 (2024).

48 Dutilh, B. E. et al. A highly abundant bacteriophage discovered in the unknown sequences of human faecal metagenomes. Nat. Commun. 5, 4498 (2014).

49 Shkoporov, A. N. et al. The human gut virome is highly diverse, stable, and individual specific. Cell Host Microbe 26, 527–541 (2019).

50 Almeida, A. et al. A unified catalog of 204,938 reference genomes from the human gut microbiome. Nat. Biotechnol. 39, 105–114 (2021).

51 Nayfach, S., Shi, Z. J., Seshadri, R., Pollard, K. S. & Kyrpides, N. C. New insights from uncultivated genomes of the global human gut microbiome. Nature 568, 505–510 (2019).

52 Hannigan, G. D., Duhaime, M. B., Ruffin IV, M. T., Koumpouras, C. C. & Schloss, P. D. Diagnostic potential and interactive dynamics of the colorectal cancer virome. mBio 9, 10.1128/mBio.02248-18 (2018).

53 Zuo, T. et al. Gut mucosal virome alterations in ulcerative colitis. Gut 68, 1169–1179 (2019).

54 Lang, S. et al. Intestinal virome signature associated with severity of nonalcoholic fatty liver disease. Gastroenterology 159, 1839–1852 (2020).

55 Yang, K. et al. Alterations in the gut virome in obesity and type 2 diabetes mellitus. Gastroenterology 161, 1257–1269 (2021).

56 Fan, G. et al. Alterations in the gut virome are associated with type 2 diabetes and diabetic nephropathy. Gut Microbes 15, 2226925 (2023).

57 Nayak, B., Liu, R. H. & Tang, J. Effect of processing on phenolic antioxidants of fruits, vegetables, and grains—a review. Crit. Rev. Food Sci. Nutr. 55, 887–918 (2015).

58 Carmody, R. N., Varady, K. & Turnbaugh, P. J. Digesting the complex metabolic effects of diet on the host and microbiome. Cell 187, 3857–3876 (2024).

59 Reyes, A. et al. Viruses in the faecal microbiota of monozygotic twins and their mothers. Nature 466, 334–338 (2010).

60 Manrique, P. et al. Healthy human gut phageome. Proc. Natl. Acad. Sci. U. S. A. 113, 10400–10405 (2016).

61 Kim, M.-S. & Bae, J.-W. Lysogeny is prevalent and widely distributed in the murine gut microbiota. ISME J. 12, 1127–1141 (2018).

62 Liang, G. et al. The stepwise assembly of the neonatal virome is modulated by breastfeeding. Nature 581, 470–474 (2020).

63 Pei, Z. et al. A universe of human gut-derived bacterial prophages: unveiling the hidden viral players in intestinal microecology. Gut Microbes 16, 2309684 (2024).

64 Kumbhari, A. et al. Discovery of disease-adapted bacterial lineages in inflammatory bowel diseases. Cell Host Microbe 32, 1147–1162 (2024).

65 Carmody, R. N. et al. Cooking shapes the structure and function of the gut microbiome. Nat. Microbiol. 4, 2052–2063 (2019).

66 Delaroque, C. et al. Diet standardization reduces intra-individual microbiome variation. Gut microbes 14, 2149047 (2022).

67 Motta, J.-P., Wallace, J. L., Buret, A. G., Deraison, C. & Vergnolle, N. Gastrointestinal biofilms in health and disease. Nat. Rev. Gastroenterol. Hepatol. 18, 314–334 (2021).

68 Rossi, E. et al. “It’s a gut feeling”–Escherichia coli biofilm formation in the gastrointestinal tract environment. Crit. Rev. Microbiol. 44, 1–30 (2018).

69 Chang, Y.-C., Lo, H.-H., Hsieh, H.-Y. & Chang, S.-M. Identification, epidemiological relatedness, and biofilm formation of clinical Chryseobacterium indologenes isolates from central Taiwan. J. Microbiol. Immunol. Infect. 48, 559–564 (2015).

70 Luis, A. S., Yates, E. A. & Cartmell, A. Functions and specificity of bacterial carbohydrate sulfatases targeting host glycans. Essays Biochem. 67, 429–442 (2023).

71 Anderson, C. L., Sullivan, M. B. & Fernando, S. C. Dietary energy drives the dynamic response of bovine rumen viral communities. Microbiome 5, 1–19 (2017).

72 de Jonge, P. A. et al. Gut virome profiling identifies a widespread bacteriophage family associated with metabolic syndrome. Nat. Commun. 13, 3594 (2022).

73 Clooney, A. G. et al. Whole-virome analysis sheds light on viral dark matter in inflammatory bowel disease. Cell Host Microbe 26, 764–778 (2019).

74 Liao, C. et al. Oral bacteria relative abundance in faeces increases due to gut microbiota depletion and is linked with patient outcomes. Nat. Microbiol. 9, 1555–1565 (2024).

75 von Hertzen, L. Plant microbiota: implications for human health. Br. J. Nutr. 114, 1531–1532 (2015).

76 Willner, D. et al. Metagenomic detection of phage-encoded platelet-binding factors in the human oral cavity. Proc. Natl. Acad. Sci. U. S. A. 108, 4547–4553 (2011).

77 Boling, L. et al. Dietary prophage inducers and antimicrobials: toward landscaping the human gut microbiome. Gut Microbes 11, 721–734 (2020).

78 Zmora, N. et al. Personalized gut mucosal colonization resistance to empiric probiotics is associated with unique host and microbiome features. Cell 174, 1388–1405 (2018).

79 Han, G., Luong, H. & Vaishnava, S. Low abundance members of the gut microbiome exhibit high immunogenicity. Gut Microbes 14, 2104086 (2022).

80 Wortelboer, K. et al. Phage-microbe dynamics after sterile faecal filtrate transplantation in individuals with metabolic syndrome: a double-blind, randomised, placebo-controlled clinical trial assessing efficacy and safety. Nat. Commun. 14, 5600 (2023).

81 Tian, X. et al. Gut virome-wide association analysis identifies cross-population viral signatures for inflammatory bowel disease. Microbiome 12, 130 (2024).

82 Bonilla, N. et al. Phage on tap–a quick and efficient protocol for the preparation of bacteriophage laboratory stocks. PeerJ 4, e2261 (2016).

83 Chen, S., Zhou, Y., Chen, Y. & Gu, J. fastp: an ultra-fast all-in-one FASTQ preprocessor. Bioinformatics 34, i884–i890 (2018).

84 Wood, D. E., Lu, J. & Langmead, B. Improved metagenomic analysis with Kraken 2. Genome Biol. 20, 1–13 (2019).

85 Nurk, S., Meleshko, D., Korobeynikov, A. & Pevzner, P. A. metaSPAdes: a new versatile metagenomic assembler. Genome Res. 27, 824–834 (2017).

86 Roux, S., Enault, F., Hurwitz, B. L. & Sullivan, M. B. VirSorter: mining viral signal from microbial genomic data. PeerJ 3, e985 (2015).

87 Ren, J., Ahlgren, N. A., Lu, Y. Y., Fuhrman, J. A. & Sun, F. VirFinder: a novel k-mer based tool for identifying viral sequences from assembled metagenomic data. Microbiome 5, 1–20 (2017).

88 Von Meijenfeldt, F. B., Arkhipova, K., Cambuy, D. D., Coutinho, F. H. & Dutilh, B. E. Robust taxonomic classification of uncharted microbial sequences and bins with CAT and BAT. Genome Biol. 20, 1–14 (2019).

89 Olm, M. R., Brown, C. T., Brooks, B. & Banfield, J. F. dRep: a tool for fast and accurate genomic comparisons that enables improved genome recovery from metagenomes through de-replication. ISME J. 11, 2864–2868 (2017).

90 Callahan, B. J. et al. DADA2: High-resolution sample inference from Illumina amplicon data. Nat. Methods 13, 581–583 (2016).

91 Rastogi, G., Tech, J. J., Coaker, G. L. & Leveau, J. H. A PCR-based toolbox for the culture-independent quantification of total bacterial abundances in plant environments. J. Microbiol. Methods 83, 127–132 (2010).

92 Větrovský, T. & Baldrian, P. The variability of the 16S rRNA gene in bacterial genomes and its consequences for bacterial community analyses. PloS One 8, e57923 (2013).

93 Hockenberry, A. J. & Wilke, C. O. BACPHLIP: predicting bacteriophage lifestyle from conserved protein domains. PeerJ 9, e11396 (2021).

94 Dion, M. B. et al. Streamlining CRISPR spacer-based bacterial host predictions to decipher the viral dark matter. Nucleic Acids Res. 49, 3127–3138 (2021).

95 Parks, D. H. et al. GTDB: an ongoing census of bacterial and archaeal diversity through a phylogenetically consistent, rank normalized and complete genome-based taxonomy. Nucleic Acids Res. 50, D785–D794 (2022).

96 Nayfach, S. et al. A genomic catalog of Earth’s microbiomes. Nat. Biotechnol. 39, 499–509 (2021).

97 Bland, C. et al. CRISPR recognition tool (CRT): a tool for automatic detection of clustered regularly interspaced palindromic repeats. BMC bioinformatics 8, 1–8 (2007).

98 Mallick, H. et al. Multivariable association discovery in population-scale meta-omics studies. PLoS Comput. Biol. 17, e1009442 (2021).

99 Yoon, S.-I. et al. Nutrient inadequacy in Korean young adults with depression: a case control study. Nutrients 15, 2195 (2023).

100 Yun, S. H., Shim, J.-S., Kweon, S. & Oh, K. Development of a food frequency questionnaire for the Korea National Health and Nutrition Examination Survey: data from the fourth Korea National Health and Nutrition Examination Survey (KNHANES IV). Korean J. Food Nutr. 46, 186–196 (2013).

